# IL-17A Mediates Cardiac Hypertrophy in Chronic Kidney Disease through the IL-23/Th17 Axis

**DOI:** 10.64898/2026.09.17.752355

**Authors:** Fumihiko Ogata, Shinsuke Hanatani, Yuki Okuno, Masahiro Yamamoto, Kei Morikawa, Takumi Nagakura, Satoshi Araki, Eiichiro Yamamoto, Yuichiro Arima, Yasuhiro Izumiya, Kenichi Tsujita

**Author notes:** Address correspondence to: Shinsuke Hanatani, MD, PhD, Department of Cardiovascular Medicine, Graduate School of Medical Sciences, Kumamoto University, Kumamoto, Japan, Address: 1-1-1 Honjo, Chuo-ku, Kumamoto 860-8556, Japan.

## Abstract

**Background:** Chronic kidney disease (CKD) is a major risk factor for cardiovascular morbidity, promoting left ventricular hypertrophy (LVH) prior to overt heart failure. The mechanisms underlying early cardiac remodeling in CKD remain incompletely understood. The IL-23/Th17/IL-17A inflammatory axis has emerged as a key mediator of cardiovascular pathology; however, its role in CKD-associated cardiac remodeling remains unclear. This study aimed to elucidate the contribution of this pathway to the early cardiac changes in CKD.

**Methods:** Using a 5/6 nephrectomy (5/6Nx) mouse model of CKD, we evaluated cardiac structure, mitochondrial function, and inflammatory signaling. Cardiomyocyte transcriptomes were analyzed by RamDA-seq to identify molecular alterations. Circulating cytokines were profiled longitudinally, and IL-17A neutralization experiments were performed to assess causality. In addition, serum IL-12p40 concentrations were measured in patients with CKD and correlated with their renal function and cardiac remodeling indices.

**Results:** At 16 weeks post-5/6Nx, mice developed LVH with preserved systolic function, accompanied by increased oxidative stress and mitochondrial structural abnormalities, but without significant myocardial fibrosis. Transcriptomic analysis identified oxidative phosphorylation as the most significantly downregulated pathway in cardiomyocytes. Despite increased mitochondrial DNA copy number and elevated PGC-1α expression, ATP production and mitochondrial respiratory chain proteins were reduced, indicating mitochondrial dysfunction. Circulating IL-12p40 and IL-23 levels increased progressively, concurrent with expansion of splenic Th17 cells and elevated cardiac IL-17A expression. Recombinant IL-17A directly impaired mitochondrial respiration in cultured cardiomyocytes. In vivo, IL-17A neutralization attenuated LVH, reduced oxidative stress, and partially restored mitochondrial protein expression without improving renal function. In patients with CKD, serum IL-12p40 levels correlated inversely with estimated glomerular filtration rate and positively with left ventricular mass index, supporting clinical relevance.

**Conclusions:** Activation of the systemic IL-23/Th17/IL-17A axis contributes to early CKD-associated cardiac remodeling by promoting oxidative stress and mitochondrial dysfunction. Circulating IL-12p40 may serve as a biomarker of cardiorenal remodeling, and the IL-23/IL-17A pathway represents a potential therapeutic target.

**CLINICAL PERSPECTIVE:** *What Is New?:*

- In a mouse model of chronic kidney disease (CKD), activation of the systemic IL-23/Th17/IL-17A axis was associated with left ventricular hypertrophy, oxidative stress, and mitochondrial dysfunction before the development of overt systolic dysfunction or myocardial fibrosis.
- In patients with CKD, circulating IL-12p40 levels were associated with both renal dysfunction and increased left ventricular mass.

*What Are the Clinical Implications?:*

- These findings identify the IL-23/Th17/IL-17A axis as a potential mechanistic link between CKD and early cardiac remodeling, suggesting that inflammation may contribute to cardiac injury independently of worsening renal function.
- Targeting the IL-23/IL-17A pathway may represent a potential therapeutic strategy to prevent or attenuate CKD-associated cardiac remodeling before the development of overt heart failure.

## Introduction

Heart failure is a growing public health challenge in aging populations, particularly in developed countries. Chronic kidney disease (CKD) has emerged as a major independent risk factor for adverse cardiovascular outcomes ^1^, including left ventricular hypertrophy (LVH) and heart failure ^2^. Cardio-renal syndrome type 4 is a pathophysiological condition in which primary CKD drives progressive cardiac dysfunction through systemic alterations, such as inflammation, metabolic disturbances, and neurohormonal activation.

In advanced CKD stages, several circulating mediators, including the uremic toxin indoxyl sulfate (IS) ^3^ ^4^, have been implicated in myocardial remodeling, as demonstrated in clinical and experimental studies. While uremic cardiomyopathy and oxidative stress have been studied extensively, these investigations have focused mainly on patients with end-stage renal disease ^5^. Epidemiological data indicate that the risk of heart failure begins to increase as early as CKD stage 3 ^6^; however, classical uremic toxins, such as p-cresol and IS, accumulate significantly only in advanced CKD or after dialysis initiation ^7^. This suggests that additional non-uremic mechanisms may contribute to early cardiac remodeling in CKD ^8,9^.

Recently, attention has shifted toward elucidating early cardiac responses in non-end-stage CKD, with emerging evidence suggesting that mitochondrial dysfunction, which is central to cellular energy metabolism, may play a key role in early CKD-induced myocardial injury ^8^. However, the precise molecular mechanisms underlying this process remain unclear.

Using a well-established 5/6 nephrectomy (5/6Nx) mouse model and serum samples from patients with CKD, we aimed to identify novel inflammatory cytokines that mediate cardio-renal interactions and drive cardiac remodeling in CKD.

## Methods

### Animal experiment

All procedures were performed in accordance with the Kumamoto University Animal Care Guidelines and the *Guide for the Care and Use of Laboratory Animals* published by the US National Institutes of Health (Publication No. 85-23, revised 1996). The experimental protocol was approved by the Animal Care and Use Committee of Kumamoto University (approval no. A2025-123). The reporting of the animal experiments followed the ARRIVE 2.0. Eight-week-old male C57BL/6J mice were purchased from Kyudo Co., Ltd. (Fukuoka, Japan) and housed at a controlled temperature of 24 °C with a 12-hour light–dark cycle. The nephrectomy and sham procedures were performed under inhalational anesthesia with 2% isoflurane. After surgery, mice were placed on a heating pad until recovery from anesthesia. General condition, activity, posture, wound condition, and signs of pain or distress were assessed everyday. Body weight (BW) was measured weekly for health monitoring, and values obtained at 4-week intervals were used for longitudinal analysis and graphical presentation. A reduction of 20% or more from preoperative body weight was defined as a humane endpoint. At the end of the study, mice were euthanized by cervical dislocation.

### Experimental design and rigor

Mice were allocated to experimental groups based on body weight to achieve comparable baseline body weights across groups. Echocardiography, quantitative outcome measurements, sample preparation for RamDA-seq, and flow cytometric acquisition and analysis were performed by investigators blinded to group allocation. Group allocation was disclosed only after completion of data acquisition and analysis.

## Data Availability

Detailed methodologies are provided in the Supplemental Methods and Key Resources Table. All supporting data, analysis methods, and study materials related to this study are available from the corresponding author upon reasonable request. The RNA-sequencing data generated in this study will be deposited in the DNA Data Bank of Japan Sequence Read Archive and made publicly available before publication.

## Results

### 5/6Nx induces left ventricular hypertrophy

At 16 weeks after 5/6Nx, a reduction in body weight was observed (Figure 1B), whereas systolic blood pressure was elevated (Figure 1C), and renal function was compromised compared with sham-operated mice (Figure 1D). Histological assessment revealed a significant increase in the glomerulosclerosis index, quantified by periodic acid–Schiff staining, in 5/6Nx mice. This was accompanied by increased interstitial fibrosis, as assessed by Sirius Red staining, indicating progressive glomerular injury and fibrotic remodeling compared with that in sham mice (Figure 1E and Figure S1A). In addition, expression of fibrosis- and inflammation-related genes in the kidneys increased significantly, as assessed by real-time PCR (Figure S1B). Echocardiography showed a preserved ejection fraction but increased left ventricular wall thickness in 5/6Nx mice (Figure 1F and Table S1). Left ventricular weight was greater in 5/6Nx mice (Figure 1G), and histological analysis confirmed an increase in cardiomyocyte cross-sectional area (Figure 1H and 1I). Consistently, the expression of LV hypertrophy–associated genes (ANP and β-MHC) was increased significantly (Figure 1J).

**Figure 1.**
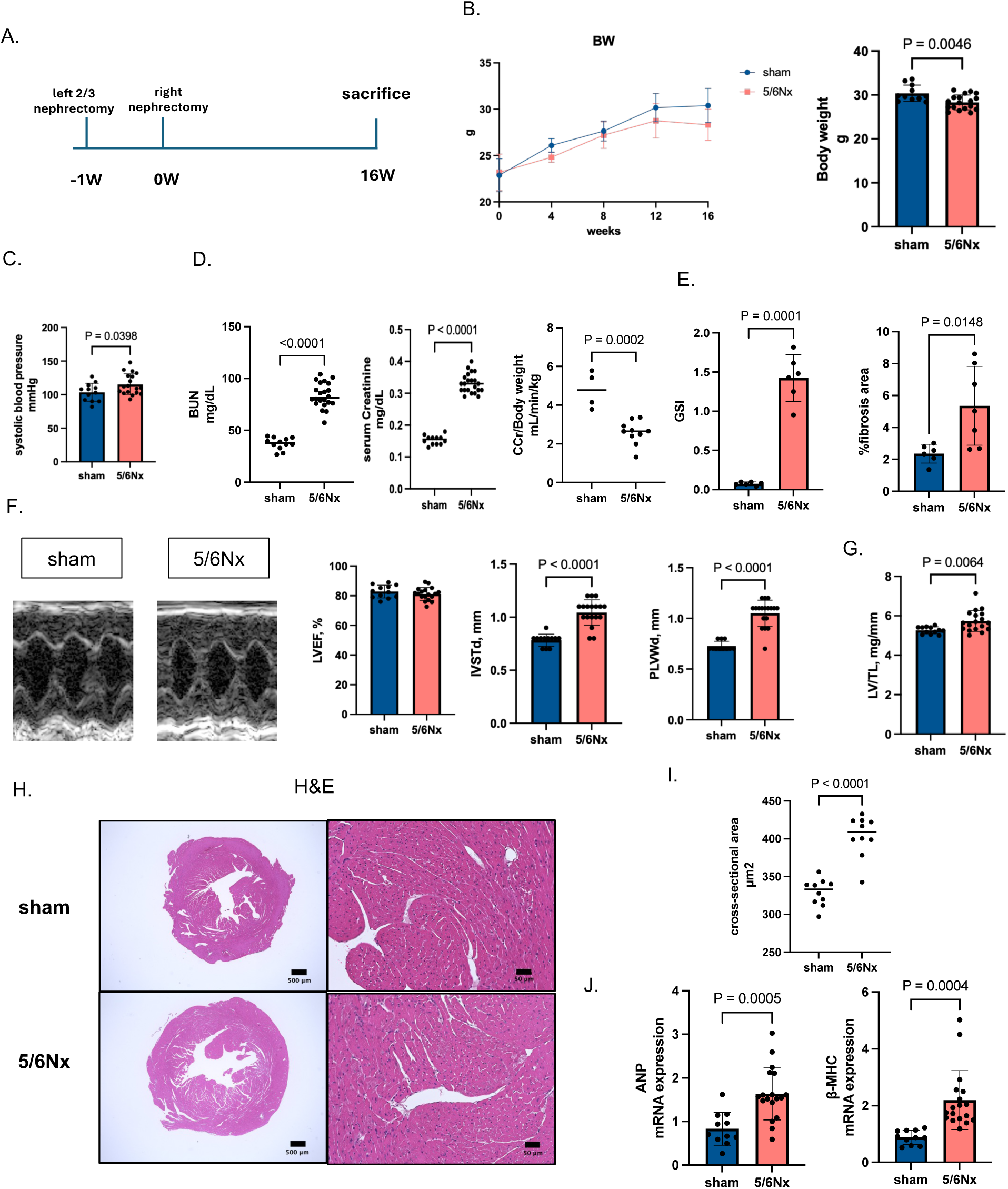
Development of cardiac hypertrophy and renal dysfunction in a 5/6 nephrectomy–induced CKD mouse model. (A) Experimental protocol for the two-step 5/6 nephrectomy (5/6Nx) model. Two-thirds of the left kidney was surgically removed, followed by right nephrectomy one week later. Mice were sacrificed 16 weeks after the initial surgery. (B) Time course of body weight (BW) changes following surgery (left) and BW at 16 weeks (right) in sham and 5/6Nx mice. Sham (n=12) vs. 5/6Nx (n=18). (C) Systolic blood pressure measured 16 weeks after surgery. Sham (n=12) vs. 5/6Nx (n=18). (D) Blood urea nitrogen (BUN) levels, serum creatinine and creatinine clearance (CCr) indicating impaired renal function in 5/6Nx mice. Sham (n = 12) and 5/6Nx (n = 18). (E) Glomerulosclerosis index (GSI; middle) and renal fibrosis area quantified by histological analysis (right). (F) Representative M-mode echocardiographic images (left) and quantitative analysis of left ventricular ejection fraction (LVEF), interventricular septal thickness (IVST), and posterior wall thickness (PLWT). Sham (n = 12) and 5/6Nx (n = 18). (G) Left ventricular mass normalized to tibial length (LV/TL). Sham (n=12) and 5/6Nx (n=18). (H) Representative hematoxylin and eosin (H&E)-stained transverse heart sections from sham and 5/6Nx mice. Scale bars: 500 μm (left) and 50 μm (right). (I) Quantification of cardiomyocyte cross-sectional area in sham and 5/6Nx mice (n = 10 per group). (J) Relative mRNA expression levels of hypertrophic markers atrial natriuretic peptide (ANP) and β–myosin heavy chain (β-MHC) in left ventricular tissue. Sham (n=11) and 5/6Nx (n=18) groups. Data are presented as mean ± SD. Each dot represents an individual mouse. Statistical significance was determined using an unpaired two-tailed Student’s *t-*test. The *P* values are indicated in each panel.

### Oxidative stress in LV tissue without overt cardiac fibrosis in 5/6Nx mice

4-HNE immunostaining was increased in 5/6Nx hearts (Figure 2A), accompanied by increased mRNA expression of NOX2 and p22phox (Figure 2B) as well as elevated protein levels of HO-1 and NQO1 (Figure 2C), indicating enhanced oxidative stress in cardiomyocytes of 5/6Nx mice. Although Sirius Red staining showed a trend toward increased interstitial fibrosis in 5/6Nx hearts, this difference was not statistically significant (Figure 2D). The mRNA expression levels of α-SMA and collagen1a1 were unchanged, whereas TGF-β1 expression was upregulated significantly (Figure 2E).

**Figure 2.**
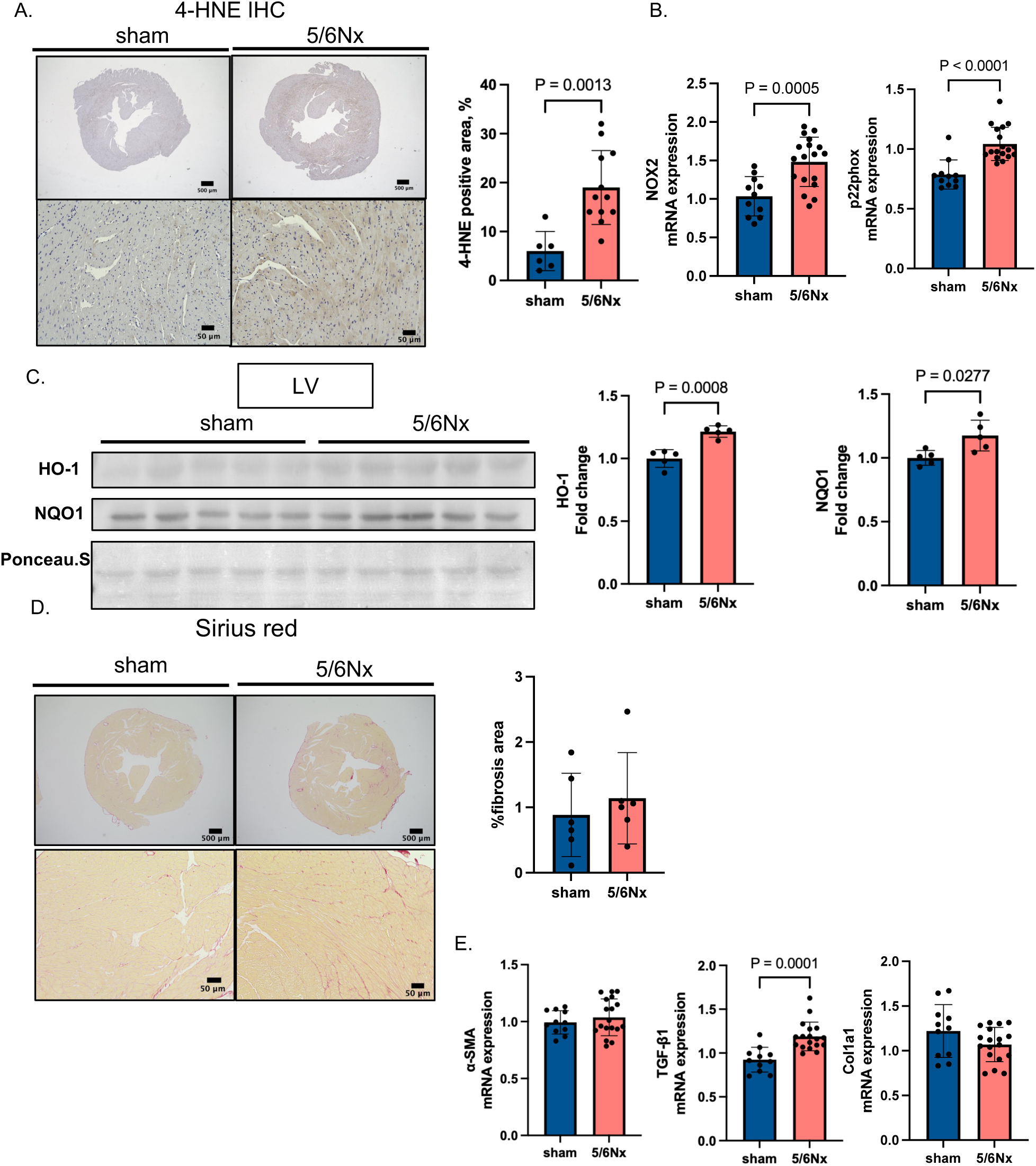
Enhanced oxidative stress in the left ventricle without overt cardiac fibrosis in 5/6 nephrectomy–induced CKD mice. (A) Representative immunohistochemical staining for 4-hydroxy-2-nonenal (4-HNE) in transverse left ventricular sections from sham and 5/6 nephrectomy (5/6Nx) mice (left) and quantitative analysis of the 4-HNE–positive area (n = 10 per group) (right). Scale bars: 500 μm (top) and 50 μm (bottom). (B) Relative mRNA expression levels of oxidative stress–related genes, including NADPH oxidase 2 (NOX2) and p22phox, in left ventricular tissue. Sham (n = 11) vs. 5/6Nx (n = 18). (C) Representative immunoblots of heme oxygenase-1 (HO-1) and NAD(P)H quinone dehydrogenase 1 (NQO1) in left ventricular lysates (n = 5 per group), with quantitative analysis normalized to staining (Ponceau S). (D) Representative Sirius Red–stained heart sections from sham and 5/6Nx mice (n = 5 per group) (left) and quantitative assessment of the fibrotic area (right). Scale bar: 500 μm. (E) Relative mRNA expression levels of fibrosis-related genes, including α-smooth muscle actin (α-SMA), transforming growth factor–β1 (TGF-β1), and collagen type I alpha 1 (Col1a1), in left ventricular tissue. Sham (n = 11) vs. 5/6Nx (n = 18). Data are presented as mean ± SD. Each dot represents an individual mouse. Statistical comparisons were performed using unpaired, two-tailed Student’s *t-*test. The *P* values are indicated in each panel.

### Oxidative phosphorylation is impaired in the hearts of 5/6Nx mice

To examine cardiomyocyte-specific transcriptional changes associated with phenotypic alterations in the hearts of 5/6Nx mice, cardiomyocytes were isolated via cell sorting and analyzed using random displacement amplification sequencing (RamDA-seq). Among the pathways with an absolute normalized enrichment score (NES) greater than 1.5, oxidative phosphorylation emerged as the most significantly downregulated pathway, exhibiting a significantly negative NES (false discovery rate < 0.05) (Figure 3A, Table S2). This was accompanied by the coordinated downregulation of genes encoding the various mitochondrial respiratory chain complexes. Heatmap visualization of variance-stabilized expression data revealed broad downregulation of oxidative phosphorylation genes (complexes I–V) in 5/6Nx hearts compared with sham hearts, indicating a global impairment of mitochondrial respiratory capacity rather than isolated complex-specific defects (Figure 3B). Transmission electron microscopy (TEM) revealed disrupted inner mitochondrial membranes and cristae in cardiomyocytes from 5/6Nx mice (Figure 3C). ATP content was reduced in 5/6Nx hearts (Figure 3D), suggesting impaired myocardial ATP production. Mitochondrial DNA (mtDNA) copy number, assessed by droplet digital PCR (ddPCR), was increased in 5/6Nx hearts (Figure 3E). Protein expression of the mitochondrial biogenesis regulator PGC-1α was elevated in 5/6Nx hearts (Figure 3F). Despite these changes, oxidative phosphorylation–related proteins ATP5A and UQCRC2 were decreased (Figure 3G). Collectively, these findings suggest mitochondrial structural disruption and impaired bioenergetic function despite the activation of compensatory mitochondrial responses in the hearts of 5/6Nx mice.

**Figure 3.**
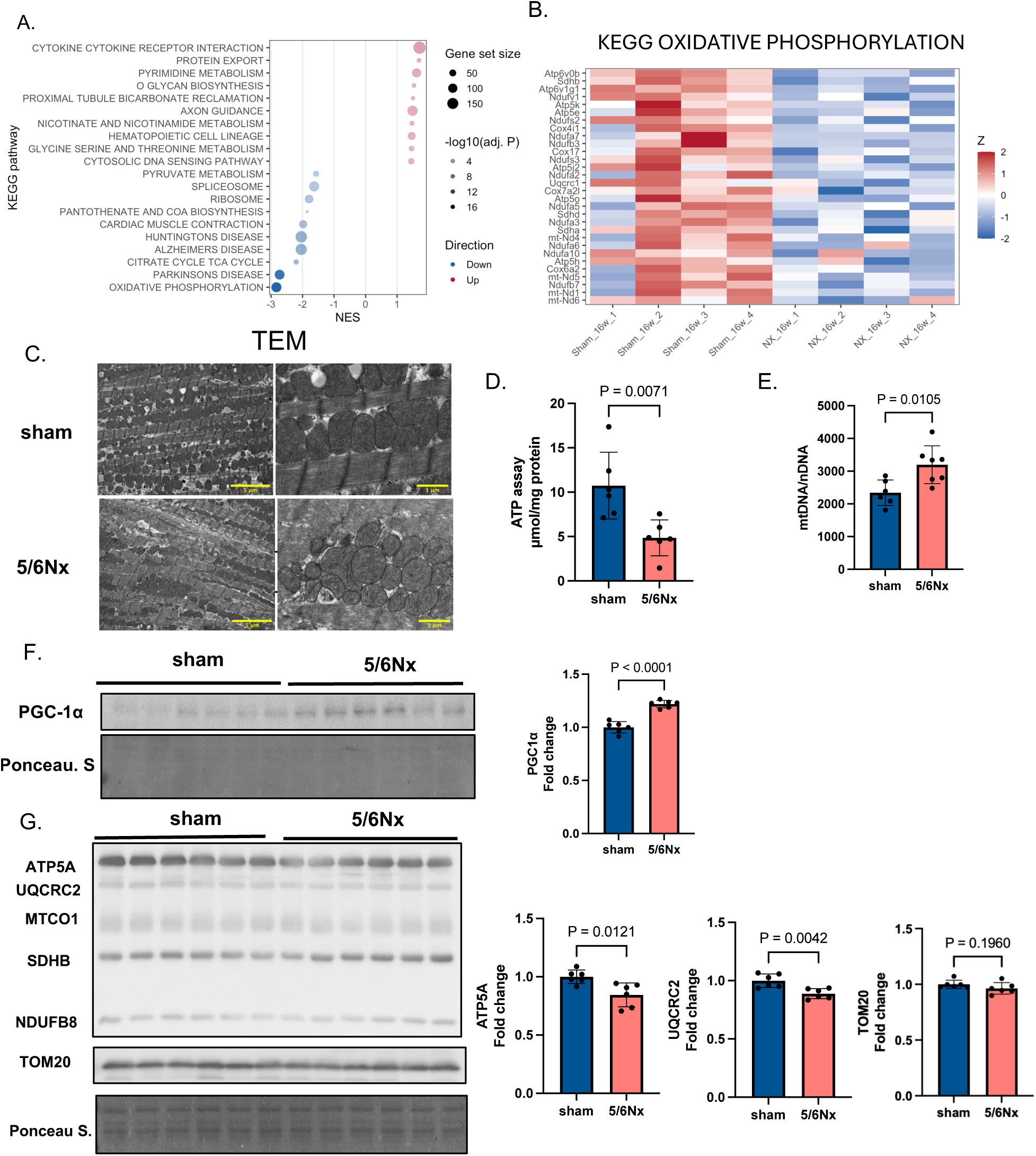
Transcriptomic profiling reveals suppression of oxidative phosphorylation–related pathways in the left ventricle of CKD mice. (A) Gene set enrichment analysis (GSEA) of KEGG pathways based on left ventricular RNA sequencing data from sham-operated and 5/6 Nx) mice. Pathways are plotted according to the normalized enrichment score (NES). Red dots indicate significantly upregulated pathways, whereas blue dots indicate downregulated pathways in 5/6Nx hearts. Dot size represents gene set size, and color intensity reflects the −log10–adjusted *P* value. (B) Heatmap of genes involved in the KEGG oxidative phosphorylation pathway. Z-score-normalized expression levels are shown for individual samples, demonstrating broad downregulation of mitochondrial electron transport chain–related genes in 5/6Nx hearts. (C) Representative transmission electron microscopy (TEM) images of left ventricular myocardium from sham and 5/6Nx mice. Mitochondria in 5/6Nx hearts exhibit disorganized cristae and structural abnormalities. Scale bars: 5 μm (left) and 1 μm (right). (D) Myocardial ATP content measured in left ventricular tissue, showing significantly reduced ATP production in 5/6Nx mice (n = 6 per group). (E) Mitochondrial DNA (mtDNA) copy number assessed by droplet digital PCR, normalized to nuclear DNA, demonstrating increased mtDNA content in 5/6Nx hearts. Sham (n =6) vs. 5/6Nx (n =7). (F) Representative immunoblots of mitochondrial biogenesis–related protein peroxisome proliferator–activated receptor γ coactivator-1α (PGC-1α) with quantitative analysis normalized to total protein staining (Ponceau S). Sham vs. 5/6Nx (n = 6 per group). (G) Representative immunoblots of mitochondrial respiratory chain–related proteins, including ATP synthase subunit alpha (ATP5A), ubiquinol–cytochrome c reductase core protein 2 (UQCRC2), mitochondrial cytochrome c oxidase subunit 1 (MTCO1), succinate dehydrogenase complex subunit B (SDHB), NADH dehydrogenase [ubiquinone] 1 beta subcomplex subunit 8 (NDUFB8), and the mitochondrial outer membrane protein TOM20 (left). Quantitative analyses normalized to total protein staining (Ponceau S) are shown in the right panel. Sham vs. 5/6Nx (n = 6 per group). Data are presented as mean ± SD. Each dot represents an individual mouse. Statistical significance was determined using an unpaired two-tailed Student’s *t-*test. The *P* values are indicated in each panel.

### IL-12p40 and IL-23 levels increase with renal decline alongside cytokine gene upregulation in 5/6Nx cardiomyocytes

We further interrogated the cardiomyocyte RamDA-seq data using unbiased pathway enrichment analysis. 5/6Nx mice exhibited a significant upregulation of genes related to cytokine–cytokine receptor interactions in the cardiomyocyte, as revealed by leading-edge gene–based pathway analysis ^10^. Notably, genes such as Cx3cl1, Epor, Cxcl14, Il2, Ifngr2, Cxcr5, Acvrl1, Ifnar2, Ifnar1, and Ifnlr1, and Il17ra were elevated compared with sham controls (Figure S2B, Table S3).

Focusing on inflammatory cytokines as mediators of organ interactions in CKD, we conducted serum cytokine profiling, which identified IL-12p40 as the cytokine increased most prominently in 5/6Nx mice (Figure 4A). ELISA confirmed a progressive rise in circulating IL-12p40 at 12 and 16 weeks post-surgery (Figure 4B). Given that IL-12p40 forms a shared subunit of the heterodimeric cytokine IL-23 and IL-12, we observed a concurrent increase in IL-23 and a decrease in IL-12 (Figure 4B). These cytokine changes occured in a time-dependent manner, correlating with worsening renal function at 16 weeks relative to 12 weeks (Figure S3A). Although left ventricular systolic function remained unchanged, left ventricular weight was increased significantly at 16 weeks compared with 12 weeks (Figure S3B).

**Figure 4.**
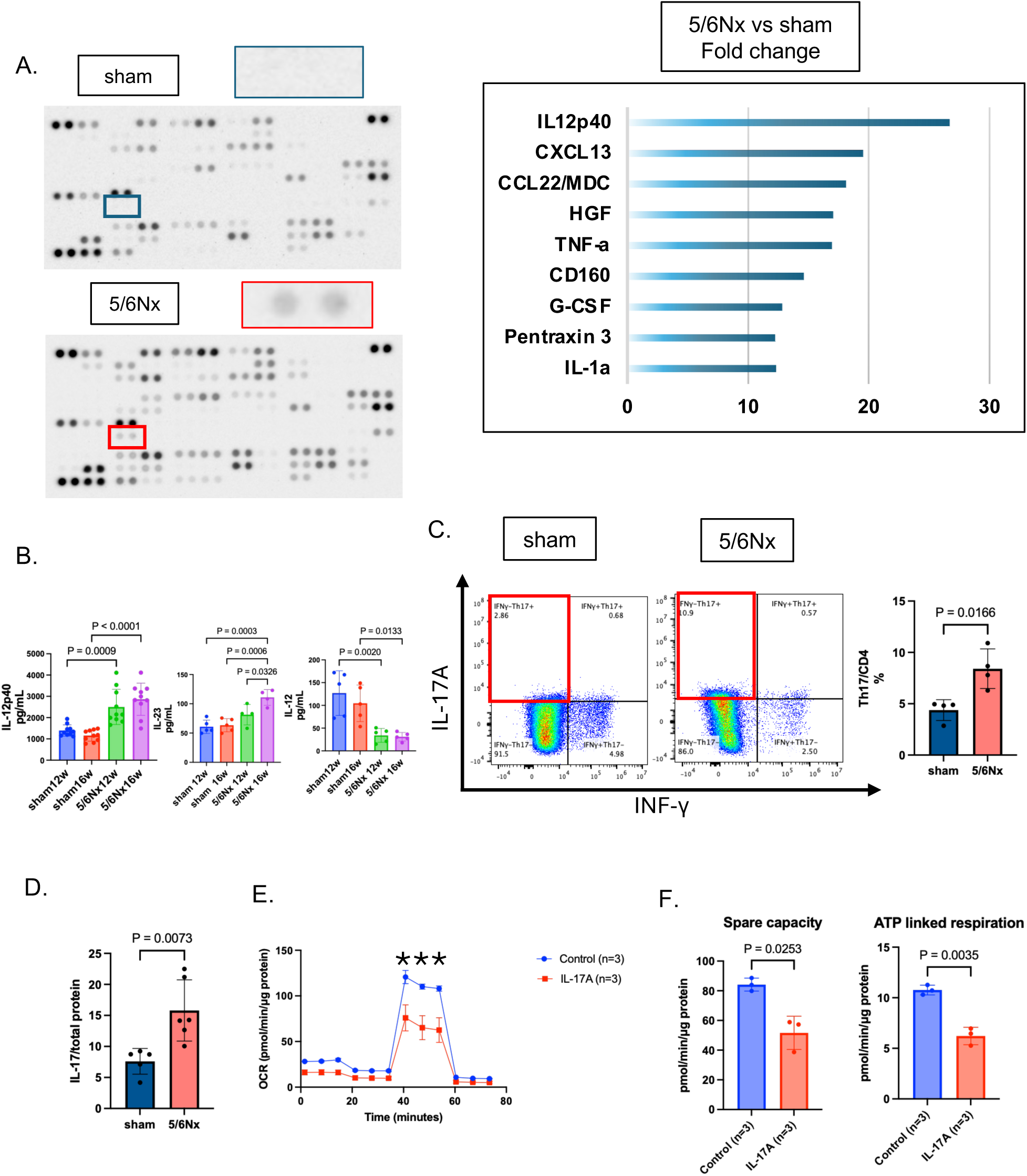
Identification of IL-12p40 as a key circulating cytokine associated with CKD. (A) Representative serum cytokine array images obtained from sham and 5/6 5/6Nx mice (left). Relative -fold changes of selected cytokines in 5/6Nx mice compared with sham controls are shown in the right panel. IL-12p40 was identified as the most prominently upregulated cytokine in CKD mice. (B) Serum concentrations of IL-12p40, measured by enzyme-linked immunosorbent assay (ELISA) at 12 and 16 weeks after surgery in sham and 5/6Nx mice (n = 10 per group). IL-23 and IL-12 levels were measured in sham and 5/6 Nx mice (n = 5 per group). (C) Representative flow cytometric analysis of splenic CD4⁺ T cells showing IL-17A and interferon-γ (IFN-γ) expression in sham and 5/6Nx mice (n = 4 per group) (left), and quantitative analysis of Th17 cells expressed as the percentage of IL-17A⁺CD4⁺ cells (right). (D) Relative protein expression levels of IL-17A in left ventricular tissue, normalized to total protein (n = 6 per group). (E) Oxygen consumption rate (OCR) measured in neonatal rat cardiomyocytes (NRCMs) using a Seahorse XF mini-analyzer after stimulation with recombinant IL-17A (50 ng/mL) or control (n = 3 per group). The time course of OCR during the mitochondrial stress test is shown in the left panel. Oligomycin, carbonyl cyanide-p-trifluoromethoxyphenylhydrazone (FCCP), and rotenone/antimycin A were injected sequentially at the indicated time. (F) Quantification of mitochondrial respiration parameters is shown in the right panels, including basal respiration, ATP-linked respiration, and spare respiratory capacity. The OCR values were normalized to protein content. Data are presented as mean ± SD (n = 3 per group). Statistical significance for the OCR time-course was determined using two-way ANOVA, and comparisons between groups for mitochondrial parameters were performed using an unpaired two-tailed Student’s t-test. *P < 0.05 vs. control. Data are presented as mean ± SD. Each dot represents an individual mouse. Statistical significance was determined using an unpaired two-tailed Student’s *t-test* or one-way analysis of variance (ANOVA) with appropriate post-hoc analysis, as indicated. The *P* values are shown in each panel.

### CKD-specific cardiac inflammation involves splenic Th17 expansion, myocardial IL-17A elevation, and mitochondrial suppression

To distinguish whether these cardiac inflammatory alterations were specific to chronic renal injury rather than reflecting acute systemic inflammation following surgery, we compared cardiac single-nucleus RNA sequencing (snRNA-seq) data from murine models of AKI and CKD ^11^. Re-analysis of these datasets demonstrated minimal Il12b (IL-12p40) expression in cardiac immune cells, indicating a scarcity of IL-12p40–and IL-23–producing cells in the heart, with only one nucleus showing detectable transcripts (Figure S4A). Consistently, Th17 transcriptional signatures and the proportion of Th17-like cells were not elevated in 5/6Nx hearts (Figure S4B and S4C). This comparison highlights that the observed cardiac inflammation is specific to CKD rather than a nonspecific postoperative response.

Since IL-23 promotes the expansion and maintenance of pathogenic Th17 cells, which are the primary source of IL-17A, we investigated Th17 responses as a potential upstream driver of IL-17A–mediated cardiac remodeling in CKD. Flow cytometric analysis of splenocytes revealed an increase in Th17 cells in 5/6Nx mice (Figure 4C; gating strategy in Figure S5). Correspondingly, IL-17A protein levels were elevated in the myocardium (Figure 4D). Functional assessment of the effect of IL-17A on cardiomyocytes showed that treatment with IL-17A suppressed mitochondrial respiration, as evidenced by a reduced oxygen consumption rate (OCR) (Figure 4E and 4F). These findings collectively support a critical role for the IL-23–Th17–IL-17A axis in mediating myocardial remodeling in CKD.

### Neutralization of IL-17A alleviates hypertrophy, oxidative stress, and mitochondrial dysfunction in 5/6Nx cardiac tissue

To elucidate the role of IL-17A in cardiac injury associated with CKD, we performed loss-of-function experiments using an anti-IL-17A neutralizing antibody in vivo (Figure 5A). Although administration of the anti-IL-17A antibody partially mitigated body weight loss (Figure 5B), it did not improve blood pressure or renal function significantly (Figure 5C and 5D). In contrast, IL-17A neutralization markedly reduced LVH compared with untreated 5/6Nx mice (Figure 5E and Table S5). Consistently, expression of NOX2 mRNA, a marker of oxidative stress, was decreased significantly following anti-IL-17A antibody treatment (Figure 5F). Furthermore, the reduced expression of mitochondrial proteins UQCRC2 and ATP5A observed in 5/6Nx hearts was partially restored by anti-IL-17A antibody treatment (Figure 5G).

**Figure 5.**
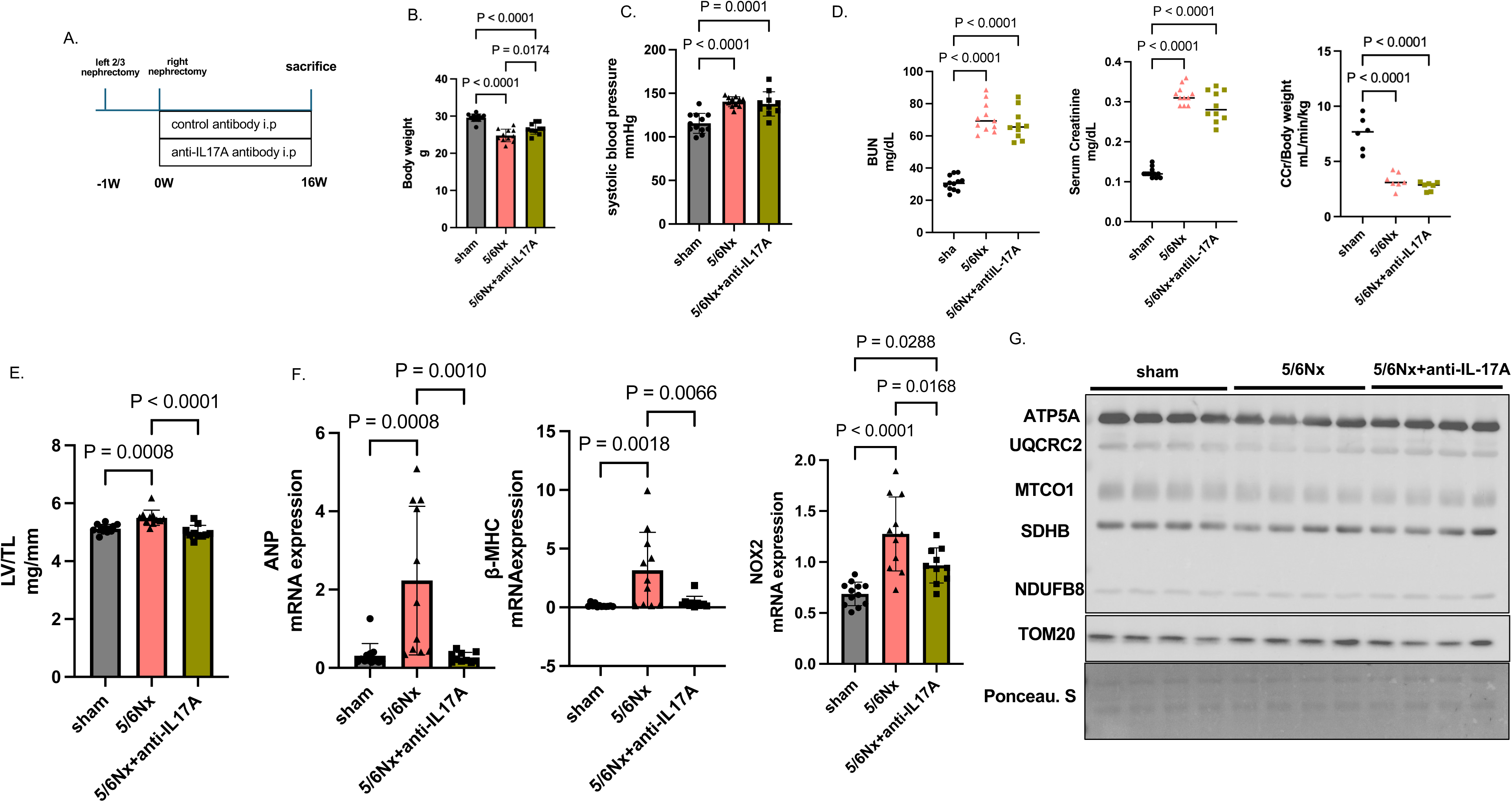
Neutralization of IL-17A attenuates cardiac hypertrophy and oxidative stress in CKD mice. (A) Experimental protocol for IL-17A neutralization in the 5/6Nx mouse model. Mice received intraperitoneal injections of control IgG or anti–IL-17A neutralizing antibody from one week after the second surgery until sacrifice at 16 weeks of age. (B) Body weight at 16 weeks in sham (n=12), 5/6Nx (n=11), and 5/6Nx mice treated with anti–IL-17A antibody (n = 10). (C) Systolic blood pressure at 16 weeks in sham (n=12), 5/6Nx (n=11), and 5/6Nx mice treated with anti–IL-17A antibody (n = 10). (D) Blood urea nitrogen (BUN), serum creatinine levels, and creatinine clearance (CCr) demonstrating partial improvement in renal dysfunction following IL-17A neutralization. BUN and serum creatinine levels were measured in sham (n = 12), 5/6Nx (n = 11), and 5/6Nx mice treated with anti–IL-17A antibody (n = 10). Creatinine clearance was measured in sham (n = 6), 5/6Nx (n = 7), and 5/6Nx mice treated with anti–IL-17A antibody (n = 7). (E) Left ventricular mass normalized to tibial length (LV/TL). Sham (n = 12), 5/6 Nx (n = 11), and 5/6Nx mice treated with anti–IL-17A antibody (n = 10). (F) Relative mRNA expression levels of atrial natriuretic peptide (ANP), β–myosin heavy chain (β-MHC) and NADPH oxidase 2 (NOX2) in left ventricular tissue. Sham (n = 12), 5/6Nx (n = 11), and 5/6Nx mice treated with anti–IL-17A antibody (n = 10). (G) Representative immunoblots of mitochondrial respiratory chain–related proteins, including ATP synthase subunit alpha (ATP5A), ubiquinol–cytochrome c reductase core protein 2 (UQCRC2), mitochondrial cytochrome c oxidase subunit 1 (MTCO1), succinate dehydrogenase complex subunit B (SDHB), NADH dehydrogenase [ubiquinone] 1 beta subcomplex subunit 8 (NDUFB8), and mitochondrial outer membrane protein TOM20 (left). Quantitative analyses normalized to total protein staining (Ponceau S) are shown in the right panel. Sham, 5/6Nx, and 5/6Nx mice treated with anti–IL-17A antibody (n = 4 per group). Data are presented as mean ± SD. Each dot represents an individual mouse. Statistical comparisons were performed using one-way analysis of variance (ANOVA), followed by Tukey’s multiple-comparison test. The *P* values are indicated in each panel.

### Clinical relevance of serum IL-12p40 in CKD-associated cardiac remodeling

Considering its clinical significance, we evaluated the serum level of IL-12p40, a component of IL-23, in patients selected according to predefined exclusion criteria (Figure S6). Compared with patients without CKD, those with CKD were older and more frequently received angiotensin receptor blocker therapy. Although the left ventricular ejection fraction did not differ significantly between groups, patients with CKD exhibited increased interventricular septal thickness at end-diastole and elevated E/e′values, an indicator of left ventricular filling pressure (Table S6). Serum IL-12p40 concentrations were significantly higher in patients with CKD with eGFR < 60 mL/min/1.73 m² and showed a negative correlation with eGFR (Figure 6A). Furthermore, serum IL-12p40 levels were elevated in patients with LVH and positively correlated with LVMi (Figure 6B). Analysis by sex reveald that this correlation between LVMi and serum IL-12p40 remained significant in both male and female CKD subgroups (Figure 6C). These findings align with murine model data supporting the role of the IL-23–Th17–IL-17A axis in CKD-specific cardiac inflammation, suggesting that myocardial IL-17A elevation and mitochondrial dysfunction are key mechanisms underlying cardiac remodeling in CKD.

**Figure 6.**
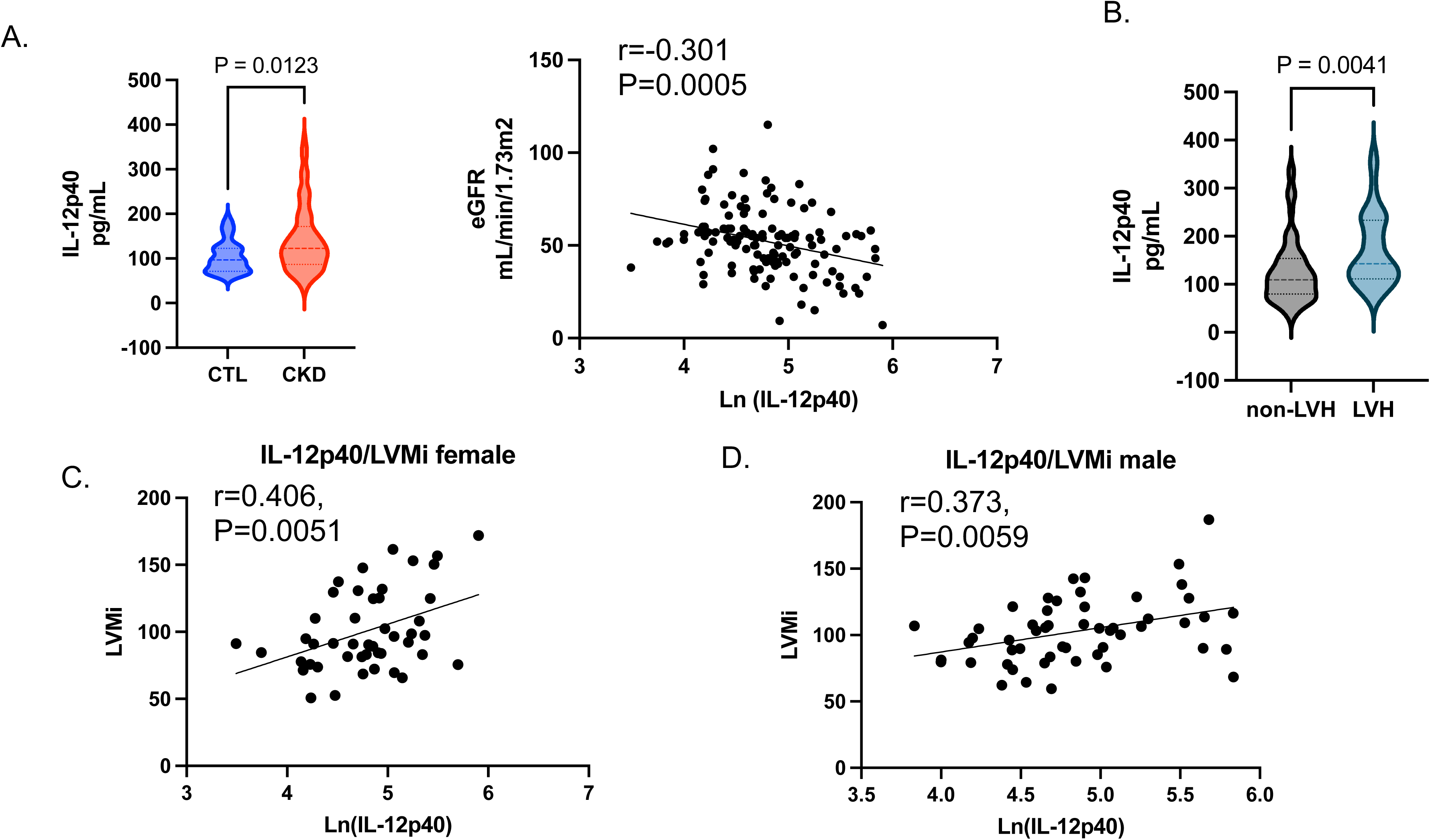
Clinical relevance of the IL-12p40 axis in patients with chronic kidney disease. (A) Serum IL-12p40 concentrations in control (CTL) patients and patients with chronic kidney disease (CKD; left) and the correlation between log-transformed IL-12p40 levels and estimated glomerular filtration rate (eGFR; right). (B) Comparison of serum IL-12p40 levels between patients with and without left ventricular hypertrophy (LVH). (C, D) Correlation between serum IL-12p40 levels and left ventricular mass index (LVMi) stratified by sex. Positive correlations were observed in both female (C) and male (D) patients. Data are presented as individual data points. Correlations were assessed using Pearson’s correlation analyses. Group comparisons were performed using unpaired two-tailed Student’s *t-*test or Mann–Whitney *U* test, as appropriate. The *P* values are indicated in each panel.

## Discussion

This study, using a CKD mouse model, demonstrated that the systemic condition of CKD precipitates LVH. The principal findings are as follows: (1) CKD induced cardiac hypertrophy accompanied by oxidative stress and mitochondrial abnormalities; (2) circulating IL-23 levels were elevated in CKD, with its downstream cytokine IL-17A promoting oxidative stress in the heart; (3) neutralization of IL-17A mitigated cardiac hypertrophy and ameliorated mitochondrial abnormalities in CKD mice; and (4) serum IL-12p40 levels correlated with renal dysfunction and LVH in patients with CKD.

In our CKD mouse model, LVH and elevated oxidative stress were observed despite the absence of overt myocardial fibrosis or systolic dysfunction. Given that mitochondrial abnormalities have been documented to precede cardiac fibrosis and dysfunction in the early stages of cardiorenal syndrome^8^, our findings suggest that this model represents the early compensatory phase of CKD-induced cardiomyopathy. Consistent with this interpretation, CKD hearts exhibited mitochondrial structural abnormalities, along with increased expression of PGC-1α and mtDNA copy number. In contrast to advanced CKD models, in which suppression of PGC-1α signaling is associated with severe cardiac hypertrophy and fibrosis^12^, these findings may reflect a compensatory response to mitochondrial stress and energy depletion during the early stages of cardiac remodeling^13^. Transcriptomic analysis revealed enrichment of genes associated with Complex I; however, these changes were not reflected at the protein level. Instead, the decreased abundance of UQCRC2 and ATP5A, along with ultrastructural abnormalities observed through transmission electron microscopy, indicated compromised mitochondrial integrity. Importantly, ATP synthase plays a critical role in maintaining the architecture of the mitochondrial cristae. Disruptions in Complex V organization have been shown to affect the structure of the inner membrane and the assembly of the respiratory chain^14,15^. Consequently, reduced ATP5A expression may signify not only a decline in ATP production, but also a disturbance in mitochondrial ultrastructural organization.

We explored the inflammatory signaling pathways linked to CKD to identify circulating mediators that may play a role in myocardial injury during the initial phases of CKD. Alterations in pathways related to inflammation and mitochondrial function were identified in purified cardiomyocytes from 5/6Nx mice. This indicates that CKD-associated inflammatory signals have a direct impact on cardiomyocytes rather than merely reflecting immune cell infiltration or interstitial remodeling. Among the circulating cytokines analyzed, IL-12p40 and IL-23 were elevated in CKD mice, potentially serving as upstream drivers of IL-17A signaling^16^. Given the pivotal role of IL-23 in maintenance and activation of pathogenic Th17 cells^17–19^, enhanced IL-23 signaling may facilitate sustained IL-17A production, leading to subsequent cardiac injury. Supporting this hypothesis, IL-17A neutralization attenuated cardiac hypertrophy in vivo. This underscores IL-17A as a critical effector and potential therapeutic target in CKD-associated cardiac remodeling. Furthermore, serum IL-12p40 levels correlated with cardiac hypertrophy in patients with CKD. Although the direct effects of IL-12p40 or IL-23 on cardiomyocyte metabolism remain unclear, elevated IL-12p40 levels have been linked to adverse cardiovascular outcomes, including heart failure, in clinical studies^20^.

Although relatively few studies have investigated the effects of IL-17A on cardiac function directly, previous research has implicated Th17/IL-17A signaling in hypertensive organ damage^21,22^. Consistent with these findings, our study identified IL-17A as a contributor to cardiac remodeling in CKD. This is the first study to reveal the pathogenic role of IL-17A in a murine model of CKD-associated cardiac injury. In a doxorubicin-induced cardiomyopathy model, excessive IL-17A induced ferroptosis via p38-p53 signaling^21^. Although GPX4, a key ferroptosis regulator^23^, was downregulated in our model, we did not detect activation of MAPK signaling (data not shown), suggesting that IL-17A may facilitate cardiac injury through alternative downstream mechanisms in CKD. In addition, systemic Th17 activation has been implicated in cardiac dysfunction associated with hyperoxalemia-related kidney disease ^24^. In our study, IL-17A was identified as a key circulating mediator of cardiac injury in 5/6Nx mice. Elevated circulating IL-17A levels have also been associated with poor prognosis in patients with heart failure. Moreover, increased IL-17A levels have been reported in individuals with CKD^25^, supporting the clinical relevance of Th17-related inflammatory signaling in CKD-associated cardiovascular disease.

Considering the potential elevation of inflammatory cytokines following the surgical 5/6Nx procedure, we assessed serum IL-12p40 levels in patients with CKD to ascertain the clinical significance of our findings. Consistent with prior studies, we observed elevated circulating IL-12p40 levels in patients with CKD, and these levels were inversely correlated with renal function. Serum IL-12p40 levels were also associated with cardiac hypertrophy, a novel finding which, to our knowledge, has not been reported previously. These results suggest a potential role for the IL-23/IL-17A axis in CKD-induced cardiac remodeling and support the translational relevance of our experimental findings to human disease. Beyond its mechanistic implications, serum IL-12p40 may serve as a clinically useful biomarker of inflammatory activity and cardiac remodeling in CKD. Moreover, therapies targeting IL-23 and IL-17A cytokines are currently available for clinical use in inflammatory disorders such as psoriasis and ulcerative colitis^26–28^. Thus, our findings provide a rationale for future translational studies targeting this pathway in CKD-associated cardiovascular diseases.

## Limitations

This study employed a model in which CKD was surgically induced; thus, the surgical procedure may have introduced stress-related effects. The human cohort consisted of a small, single-center sample from our institution, with reasons for hospitalization including ischemic heart disease, arrhythmia, peripheral artery disease, and heart failure. Therefore, further studies are warranted to evaluate circulating IL-12p40 levels in patients with CKD without concurrent cardiovascular disease requiring hospitalization.

## Conclusion

Activation of the systemic IL-23/Th17/IL-17A pathway contributes to the development of LVH associated with CKD by inducing oxidative stress and mitochondrial dysfunction in cardiomyocytes. Neutralizing IL-17A attenuated cardiac remodeling, and elevated circulating IL-12p40 levels were associated with both kidney dysfunction and increased left ventricular mass, indicating its potential as a biomarker for cardiorenal syndrome.

## Acknowledgments

We thank Megumi Nagahiro and Saeko Tokunaga for their excellent technical assistance. We also thank Dr. Takahisa Nakamura (Department of Hematology, Kumamoto University Hospital) for his assistance with the flow cytometry experiments.

## Supplemental Material

Supplemental Methods

Key Resources Table

Tables S1–S5

Unedited membrane and gels

Figure S1-6

References 29–48

