## Supplemental Data 1 for "IL-17A Mediates Cardiac Hypertrophy in Chronic Kidney Disease through the IL-23/Th17 Axis"

**Title:**

**Running title:**

**IL-17A–Mediated Cardiac Hypertrophy in CKD**

#### Supplemental Methods

##### *CKD mouse model*

A mouse model of chronic kidney disease (CKD) model was established using a two-step 5/6 nephrectomy (5/6Nx) procedure in 8-week-old C57BL/6J mice, as previously described<sup>29, 30</sup>. Briefly, a two-thirds nephrectomy of the left kidney was performed, followed by total nephrectomy of the right kidney one week later (n = 18). The sham-operated control group (n = 12) underwent all surgical steps except nephrectomy, including blunt dissection of the renal capsule and adrenal gland. Mice were maintained for 16 weeks after the final surgery. The experimental unit was an individual animal. An independent cohort of 5/6Nx (n = 14) and sham-operated mice (n = 12) was maintained for 12 weeks after the final surgery for additional experiments.

##### *Measurement of systolic blood pressure and echocardiography*

Systolic blood pressure was measured using a tail-cuff blood pressure analyzer (Muromachi, Tokyo, Japan) before sacrifice. Echocardiographic assessment of left ventricular (LV) function was performed at the same time point under awake conditions using the Xario system (Toshiba, Tokyo, Japan). Echocardiography was conducted by a physician blinded to the mouse groups. LV wall thickness, LV end-systolic and end-diastolic dimensions, and LV ejection fraction (LVEF) were measured using M-mode.

*Serum and urine collection and analysis*

Renal function was assessed through serum and urine analyses. Blood was collected via cardiac puncture at sacrifice. Serum creatinine levels were measured using a JCA-BM6050 analyzer. Urine samples were collected from metabolic cages three days prior to sacrifice. Urinary creatinine was measured using the turbidimetric immunoassay (TIA) method at SRL, Tokyo, Japan. Creatinine clearance (CCr) was calculated using the standard formula based on urine and plasma creatinine concentrations. CCr normalized to body weight (ml/min/g) was calculated as follows: $CCr = (U_{Cr} \times 24h$ $volume) / (P_{Cr} \times min) / \text{body weight}$.

*Histological staining*

Heart and kidney tissues were fixed in 4% paraformaldehyde (PFA) at 4°C for 24 hours, then transferred to 70% ethanol for storage. Samples were embedded in paraffin and sectioned at 10 µm thickness. Heart sections were stained with hematoxylin and eosin (H&E) and Sirius Red, while kidney sections were stained with periodic acid–Schiff (PAS) and Sirius Red. Histological images were acquired using an all-in-one fluorescence microscope (Keyence BZ-X700). Cardiomyocyte cross-sectional area was quantified with ImageJ software (NIH), measuring approximately 10 cardiomyocytes per LV tissue section. The glomerulosclerosis index (GSI) was evaluated according to previously published methods<sup>31, 32</sup>. Briefly, 10 to 20 glomeruli per kidney sample were assessed. The extent of glomerular sclerosis was determined by examining all glomeruli in a kidney cross-section and calculating the percentage involved. A semiquantitative score (GSI) graded sclerosis severity per glomerulus from 0 to 4+ as follows: 0 represents no lesion, 1+ represents sclerosis of <25% of the glomerulus, while 2+, 3+, and 4+ represent sclerosis of 25% to 50%, >50% to 75%, and >75% of the glomerulus. The mean GSI was calculated for each specimen. Fibrotic areas in cardiac and renal tissues were quantified using ImageJ software. Six to eight randomly selected microscopic fields per sample were analyzed, and the average percentage of fibrotic area was reported.

###### *Immunohistochemical staining*

Paraffin-embedded heart sections were deparaffinized and treated with 3% hydrogen peroxide to block endogenous peroxidase activity, followed by antigen retrieval using citrate buffer (pH 6.0) with microwave heating. Sections were then blocked with 5% goat serum for 1 hour and incubated overnight at 4°C with anti-4-HNE antibody. Subsequently, samples were incubated with an HRP-conjugated anti-rabbit secondary antibody, and visualization was performed using DAB for approximately 2 minutes. Images were acquired using an all-in-one microscope (Keyence BZ-X700), and positively stained areas were quantified using BZ-X Analyzer software (Keyence). Antibodies used are listed in the Key Resources Table.

###### *Transmission electron microscopy*

Transmission electron microscopy (TEM) was performed to evaluate mitochondrial ultrastructure in

cardiac tissue. Left ventricular samples were excised and immediately processed for TEM analysis at Tokai Electron Microscopy, Inc. (Aichi, Japan) using standard fixation and embedding protocols. Cardiac tissues from one sham mouse and one 5/6Nx mouse were fixed in 2% paraformaldehyde and 2% glutaraldehyde in 0.1 M cacodylate buffer (pH 7.4) at 4°C overnight. After three washes with 0.1 M cacodylate buffer for 30 min each at 4°C, samples were post-fixed with 2% osmium tetroxide (OsO<sub>4</sub>) in 0.1 M cacodylate buffer at 4°C for 2 hours. Samples were dehydrated through graded ethanol series (50% ethanol at 4°C for 30 minutes, 70% ethanol at 4°C for 30 minutes, 90% ethanol at room temperature for 30 minutes, and absolute ethanol at room temperature for 30 minutes, repeated three times), followed by overnight incubation in absolute ethanol at room temperature. Samples were infiltrated twice with propylene oxide (PO) for 30 minutes at room temperature, then incubated for 1 hour with a 7:3 mixture of PO and Quetol-812 resin (Nisshin EM Co., Tokyo, Japan) at room temperature. After overnight resin infiltration at room temperature, tissues were embedded in Quetol-812 resin and polymerized at 60°C for 48 hours. Ultrathin sections (70 nm) were prepared using an ultramicrotome (Ultracut UCT; Leica, Vienna, Austria) equipped with a diamond knife and mounted on copper grids. Sections were stained with 2% uranyl acetate for 15 minutes and lead stain solution (Sigma-Aldrich Co.) for 3 minutes at room temperature. Images were acquired using a transmission electron microscope (JEM-1400Plus; JEOL Ltd., Tokyo, Japan) operated at 100 kV acceleration voltage and equipped with a CCD camera (EM-14830RUBY2; JEOL Ltd.).

*Cardiomyocyte Nuclear Isolation*

Heart tissue samples (20–40 mg) were obtained from sham mice (n = 4) and 5/6Nx mice (n = 4). Cardiomyocyte nuclear isolation was performed as previously described <sup>33</sup>. The tissue was finely minced using a razor blade and transferred to a Potter-type homogenizer. The pestle was positioned just below the liquid surface and moved up and down five times at 1,000 rpm to homogenize the tissue. The homogenate was then mixed with Percoll<sup>®</sup> density gradient medium (Cytiva, Marlborough, MA, USA; Cat. No. 17089101), and the final Percoll concentration was adjusted to 27%. Samples were layered into ultracentrifuge tubes in the following order from top to bottom: 12%, 27% (sample), 31%, and 35% Percoll solutions. The tubes were centrifuged at 24,000 rpm for 10 minutes. Nuclear extracts were collected from the bottom layer and used for subsequent experiments. Cell sorting and flow cytometry analyses were performed using a MoFlo XDP cell sorter (Beckman Coulter, Brea CA, USA). Nuclei were stained with Alexa Fluor 488–conjugated anti-lamin A/C antibody (Cell Signaling Technology, Danvers, MA, USA; #8617) and Alexa Fluor 647–conjugated anti-PCM1 antibody (Santa Cruz Biotechnology, Dallas, TX, USA; sc-398365 AF647). After staining, fluorescence intensity was measured using a flow cytometer.

*Random displacement amplification sequencing analysis*

The random displacement amplification sequencing (RamDA-seq) protocol was modified for nuclei as described by Hayashi et al.<sup>34</sup>. After 1,000 nuclei were sorted, RamDA-seq reactions were performed to generate cDNA libraries using the NEBNext Ultra DNA Library Prep Kit for Illumina (New England Biolabs, Ipswich, MA, USA). Libraries were sequenced on a NovaSeq platform (Illumina, San Diego, CA, USA) using 150-bp paired-end reads. The resulting FASTQ files were processed for adaptor trimming and quality control using Trimmomatic and FastQC. Transcript quantification was performed using Salmon software (https://combine-lab.github.io/salmon/).

###### *RNA-seq data processing and differential expression analysis*

Transcript-level quantification was performed using Salmon software. Gene-level expression matrices were generated by aggregating transcript counts based on the Ensembl mouse annotation (Ensembl.Mmusculus.v79) using the AnnotationDbi package in R. Transcripts were mapped to the corresponding Ensembl gene IDs and gene symbols, and expression values were summarized at the gene level. The resulting gene-level expression matrix was used for downstream analysis, including differential expression analysis, gene set enrichment analysis, and heatmap visualization. Differential expression analysis was performed using the DESeq2 package (version 1.44.0) in R (version 4.4.2). A DESeqDataSet object was constructed from the gene-level count matrix and sample metadata, and normalization was performed using the median-of-ratios method implemented in the DESeq2. Gene-wise dispersions were estimated and fitted using the default model, and differential expression between sham and 5/6Nx groups was assessed using the Wald test. P-values were adjusted for multiple testing using the Benjamini–Hochberg false discovery rate (FDR) correction.

*Gene set enrichment analysis*

Gene set enrichment analysis (GSEA) was conducted using the fgsea package. Genes were ranked based on the signed Wald statistic obtained from DESeq2, which preserves both the direction and magnitude of differential expression. KEGG pathway gene sets were retrieved from the Molecular Signatures Database (MSigDB; collection C2: CP: KEGG). For each pathway, normalized enrichment scores (NES), adjusted *P*-values (FDR), and gene set sizes were computed using the fgsea multilevel algorithm. Pathways with an adjusted *P* value < 0.25 were considered enriched, in accordance with standard GSEA recommendations<sup>35</sup>. Enriched pathways were classified as upregulated (NES > 0) or downregulated (NES < 0), based on the sign of the NES. For visualization, the top pathways were selected according to their absolute NES values. Dot plots were generated to represent these enriched pathways, where the x-axis corresponds to NES, the size of each dot reflects the gene set size, and the intensity of the dot color indicates the $-\log_{10}$ of the adjusted *P* value. Pathways enriched in the 5/6Nx and sham groups were distinguished by color coding.

*Heat map construction of pathway-associated genes*

Leading-edge genes were extracted from the fgsea results for significantly enriched KEGG pathways. For visualization, the top 20 leading-edge genes contributing most strongly to the enrichment of each pathway were selected. Variance-stabilizing transformation (VST) was applied to the count data using

DESeq2, followed by gene-wise Z-score transformation across samples<sup>36</sup>. Heat maps were generated using the pheatmap package, applying hierarchical clustering to the leading-edge genes while maintaining a fixed sample order based on experimental groups. A fixed color scale (Z score, -2 to +2) was applied uniformly across all heat maps.

###### *Pathway score calculation*

To assess pathway activity quantitatively at the sample level, pathway scores were calculated using all leading-edge genes. For each sample, variance-stabilized expression values of the leading-edge genes were normalized by gene-wise Z-score transformation. The pathway score for each pathway was defined as the mean Z-score of all leading-edge genes<sup>37</sup>. This approach yields a relative pathway activation score, where higher values indicate increased expression of pathway-associated genes compared with the overall sample distribution. Separate pathway scores were calculated for oxidative phosphorylation-related and cytokine-related pathways<sup>37</sup>. Scatter plots were generated to examine relationships between oxidative phosphorylation and cytokine pathway scores across individual samples. Spearman's rank correlation coefficient was used to evaluate associations between pathway scores, considering the small sample size and non-parametric data distribution. Correlation coefficients ($\rho$) and corresponding *P*-values were calculated using the cor.test function in R. Scatter plots display individual samples colored by experimental group, with correlation statistics presented within the figure. The regression lines were omitted to avoid overinterpretation in the small sample analyses.

*Cytokine array*

Serum cytokine profiles in CKD mice were assessed using a cytokine array. Pooled serum samples from four 5/6Nx mice were used for the assay. The cytokine array was performed following the manufacturer’s instructions (Key Resources Table). Cytokine signal intensities were normalized to the reference spots provided in the kit, as specified in the protocol. Array images were analyzed using the ImageJ software.

*Enzyme-linked immunosorbent assay*

Serum IL-12p40, IL-12p19, IL-12p35, and IL-17A levels in blood samples collected from mice euthanized 16 weeks after 5/6Nx or sham surgery were quantified using commercially available enzyme-linked immunosorbent assay (ELISA) kits. Serum IL-12p40, IL-12p19, and IL-12p35 concentrations were measured using ELISA kits according to the manufacturer’s instructions. For IL-17A measurement, LV tissue protein lysates were used. IL-17A levels in cardiac tissue were normalized to total protein content, as previously described<sup>38</sup>. All ELISA procedures were performed according to the manufacturer’s protocols (Key Resources Table).

*Public Single-nucleus RNA-seq dataset analysis*

Single-nucleus RNA sequencing (snRNA-seq) data from a murine CKD model (GSE180852)<sup>11</sup> were analyzed using Seurat (v5). Immune cells were identified based on canonical immune markers and subsequently re-analyzed. Il12b expression was assessed using normalized RNA counts, and nuclei with Il12b expression greater than zero were classified as Il12b-positive. To evaluate Th17 transcriptional activity, a Th17 module score (Th17Score1) was calculated using the Seurat function AddModuleScore, based on canonical Th17-related genes (Rorc, Ccr6, Il23r, Rora, Stat3, Batf, and Irf4).

###### *Neutralizing antibody treatment*

An anti-IL-17A antibody was administered intraperitoneally to 5/6Nx mice (n = 12) at a dose of 50 µg per mouse twice weekly<sup>39,40</sup>. An isotype control antibody was administered to a separate group of 5/6Nx mice (n = 12) in parallel. Antibody treatment was initiated immediately after 5/6Nx surgery and continued for 16 weeks. Sham-operated mice receiving isotype control antibodies were included as controls (n = 12). At the end of the study period, mice were sacrificed, and tissue samples were collected for further analyses. Details of the antibodies used are provided in the Key Resources Table. Twelve mice were initially allocated to each group. One mouse in the 5/6Nx plus isotype-control group and two mice in the 5/6Nx plus anti-IL-17A group died during the postoperative period and were excluded from subsequent analyses. No additional animals were excluded from the analyses.

*Flow cytometry of the spleen*

Mouse spleens were harvested and immediately passed through a 70-μm cell strainer to prepare a single-cell suspension. Red blood cells were lysed using ACK buffer, and the reaction was quenched with PBS. Cells were centrifuged at $400 \times g$ for 5 minutes, and the pellet was resuspended in RPMI-1640 supplemented with GlutaMAX, 10% fetal bovine serum (FBS), 1 mM non-essential amino acids, 1 mM sodium pyruvate, 100 U/mL penicillin, and 100 μg/mL streptomycin. Cells were adjusted to a concentration of $2.0 \times 10^6$ cells/mL and stimulated with a Cell Activation Cocktail (with Brefeldin A; BioLegend) at 2 μL/mL. After 6 hours of incubation, cells were stained with a Live/Dead viability dye at room temperature, followed by surface staining with antibodies against CD45, CD3, and CD4. Following fixation and permeabilization with intracellular fixation buffer (BioLegend), cells were stained with antibodies against IL-17A and interferon-γ (IFN-γ). Th17 cells were detected using a Novocyte Penton (Agilent), and their percentage was analyzed using FlowJo version 10.8. Gates for IL-17A and IFN-γ were set using isotype controls (Figure S4). Detailed information on the antibodies is provided in the Key Resources Table.

*Droplet digital PCR*

To quantify mitochondrial copy number, a droplet digital PCR (ddPCR) assay was performed following a previously described protocol<sup>41</sup>. Total DNA was extracted from left ventricular tissue samples using the QIAamp DNA Mini kit (Qiagen) according to the manufacturer's instructions. DNA

concentrations were measured using a NanoDrop™ 2000 Spectrophotometer (Thermo Fisher Scientific). Templates of 100 ng and 0.1 ng of DNA were prepared for genomic and mitochondrial DNA, respectively. Each PCR reaction contained 10 µL 2 × ddPCR supermix for probes without UTP (Bio-Rad), 1 µL 20 × PrimeTime standard qPCR assay (Integrated DNA Technologies), template DNA, and nuclease-free water to a final volume of 20 µL. Droplets were generated using a QX200 automated droplet generator (Bio-Rad). Amplification was carried out under the following cycling conditions: 95 °C for 10 minutes; 40 cycles of 94 °C for 30 seconds followed by 60 °C for 60 seconds; 98 °C for 10 minutes; and 4 °C hold. Droplets were analyzed immediately using a QX200 droplet reader (Bio-Rad). Specific ND1 primers were used for mitochondrial DNA, and specific Ago1 primers were used for genomic DNA. Primer sequences are provided in the Key Resources Table.

###### *ATP Measurement in Cardiac Tissue*

ATP levels in cardiac tissue were measured using an alkaline extraction method followed by a luciferase-based assay (Cat. No. 346-09793; FUJIFILM Wako Pure Chemical Corporation). Mouse hearts were excised and perfused with cold saline to remove residual blood before analysis. The tissue samples were weighed and processed on ice. Cardiac tissue (e.g., 20 mg) was homogenized in ice-cold 0.5 mol/L KOH (5 µL/mg tissue; e.g., 100 µL for 20 mg) using a Dounce homogenizer. The homogenization vessel was rinsed with an equal volume of 0.5 mol/L KOH, and the wash was combined with the original homogenate. Subsequently, distilled water (twice the initial KOH volume;

e.g., 200 μL) was added, mixed thoroughly, and incubated on ice for 5 minutes. Samples were centrifuged at $12,000 \times g$ for 5 minutes at $4^{\circ}\text{C}$, and the supernatant was collected. An aliquot (e.g., equivalent to 900 μL) was neutralized with 1 mol/L $\text{KH}_2\text{PO}_4$ (approximately 0.2 times the volume; e.g., 40 μL for 200 μL extract), mixed, and incubated on ice for 5 minutes. After a second centrifugation ($12,000 \times g$, 5 minutes, $4^{\circ}\text{C}$), the supernatant was collected as the ATP measurement sample. ATP concentrations were determined using a standard curve (0–2.5 $\mu\text{mol/L}$) prepared in a dilution buffer consisting of 0.5 mol/L KOH and 1 mol/L $\text{KH}_2\text{PO}_4$ (9:5). Luminescence was measured using a microplate reader, and ATP levels were calculated using the standard curve. To account for differences in tissue input, ATP levels were normalized to the total protein content measured from parallel samples using a bicinchoninic acid assay and expressed as nmol ATP per mg protein.

###### *Mitochondrial respiration assay*

Mitochondrial respiration was assessed using a Seahorse XF HS Mini Analyzer (Agilent Technologies). One day before the assay, the sensor cartridge was hydrated overnight in XF calibrant at $37^{\circ}\text{C}$ in a $\text{CO}_2$-free incubator. NRCMs were isolated from 2-day-old Wistar rats purchased from Japan SLC, Inc. (Shizuoka, Japan). Sex was not determined because of the neonatal age. Cardiomyocytes isolated from multiple pups were pooled for each cell preparation, as described previously<sup>42</sup>. Cells were seeded at a density of $1.0 \times 10^4$ cells per well in Seahorse XF microplates and cultured in low-glucose DMEM supplemented with 10% fetal bovine serum (FBS) for 2 days. The cells were then serum-starved in

low-glucose DMEM containing 1% FBS overnight. Recombinant IL-17A (50 ng/mL) was added to stimulate the cells for 12 hours. On the day of the assay, the cells were washed, and the culture medium was replaced with Seahorse XF assay medium consisting of XF Base Medium supplemented with 2 mM glucose, 1 mM sodium pyruvate, and 2 mM glutamine. Cells were incubated in 150 μL of assay medium for 1 hour at 37 °C in a CO<sub>2</sub>-free incubator to allow temperature and pH equilibration. For the mitochondrial stress test, oligomycin (1 μM), carbonyl cyanide-p-trifluoromethoxy phenylhydrazone (FCCP, 1 μM), and a mixture of antimycin A (0.5 μM) and rotenone (0.5 μM) were injected sequentially through the ports of the hydrated sensor cartridge. The oxygen consumption rate (OCR) was measured under basal conditions, followed by sequential injections of oligomycin, FCCP, and antimycin A/rotenone. Basal respiration, ATP-linked respiration, maximal respiration, and spare respiratory capacity were calculated using Seahorse XF software. OCR values were normalized to total cellular protein content measured using a bicinchoninic acid (BCA) protein assay. Each condition was measured in triplicate, and experiments were independently repeated at least three times.

###### *Quantitative real-time PCR*

Messenger RNA (mRNA expression in LV and kidney tissues) was evaluated using quantitative real-time polymerase chain reaction (qRT-PCR). The protocol was described previously <sup>43</sup>. Primers used are listed in the Key Resources Table.

*Western Blot Analysis*

Western blot analysis was performed using an SDS-PAGE system, as described previously <sup>44</sup>. Details of the antibodies used are provided in Key Resources Table.

*CKD population*

Patients with CKD comprised a consecutive series hospitalized for cardiac disease between January and December 2019 in the Department of Cardiovascular Medicine at Kumamoto University Hospital. Patient characteristics are summarized in Table S6. Initially, 1,439 patients hospitalized during this time period were screened. Serum IL-12p40 levels were measured in 99 patients after applying the following exclusion criteria: preserved renal function (eGFR > 60 mL/min/1.73 m<sup>2</sup>; n = 705), hemodialysis treatment (n = 91), active malignant tumors (n = 26), and history of nephrectomy (n = 2). Additional exclusions were conditions affecting cardiac structure or systemic inflammation: severe valvular heart disease (severe aortic stenosis, n = 52; severe aortic regurgitation, n = 8; severe mitral stenosis, n = 6; severe mitral regurgitation, n = 13), infective endocarditis (n = 2), pericarditis (n = 3), cardiomyopathy (n = 82), vasculitis (n = 4), aortic aneurysm (n = 56), pulmonary hypertension (n = 22), connective tissue diseases (rheumatoid arthritis <sup>45,46</sup>, n = 3; systemic lupus erythematosus, n = 1; systemic sclerosis, n = 1), and inflammatory bowel disease <sup>47</sup> (n = 3). Patients admitted on an emergency basis (n = 164) and those without cardiac catheterization (n = 96) were also excluded. For the control group, consecutive patients admitted in January 2019 with preserved renal function (eGFR

> 60 mL/min/1.73 m<sup>2</sup>) were screened. Applying the same exclusion criteria as for the CKD group, 26 patients were selected as controls. Transthoracic echocardiography was performed upon hospital admission. Body weight and height were measured at the time of echocardiography, and the left ventricular mass index (LVMI) was calculated by indexing left ventricular mass to body surface area. Left ventricular mass was calculated using the Devereux formula based on linear echocardiographic measurements<sup>48</sup>.

#### *Statistical Analysis*

All in vivo and in vitro experimental data are presented as mean ± standard deviation (SD). Statistical analyses were performed using GraphPad Prism (version 8.02; GraphPad Software, La Jolla, CA, USA). Data distributions were assessed using the Shapiro-Wilk test, and homogeneity of variance was evaluated using the Brown-Forsythe test. Comparisons between two groups were performed using an unpaired Student's t-test for normally distributed data with equal variances, Welch's t-test for normally distributed data with unequal variances, or the Mann-Whitney U test for non-normally distributed data. Comparisons among multiple groups were performed using ordinary one-way ANOVA followed by Tukey's multiple-comparisons test when parametric assumptions were met. When these assumptions were not met, Welch's ANOVA or the Kruskal-Wallis test followed by an appropriate multiple-comparisons test was used. Time-course OCR data obtained from Seahorse assays were analyzed using two-way repeated-measures ANOVA with treatment group and time as factors, followed by Sidak's multiple comparisons test, where appropriate. Correlations between continuous variables were assessed using Pearson's correlation coefficient. All statistical analyses related to RNA sequencing and gene set enrichment analysis (GSEA) were performed using the R software. Data are presented as individual data points, unless otherwise stated. Statistical significance was defined as a two-sided *P*-value $< 0.05$, unless otherwise specified. For human clinical data, serum high-sensitivity troponin T (hs-TnT), B-type natriuretic peptide (BNP), and interleukin-12p40 (IL-12p40) levels were analyzed using nonparametric methods because of their non-normal distributions and are presented as medians with interquartile ranges (IQR). Other continuous variables with normal distributions are presented as mean ± SD and were compared using Student's t-test. Categorical variables were compared using the chi-square test. Graphical presentations were generated using GraphPad Prism, and statistical analyses of human clinical data were performed using SPSS Statistics version 27 (IBM Corp., Armonk, NY, USA).

1
2
3

##### Key Resources Table

| Mouse primers for qPCR |  |  |
| --- | --- | --- |
| Gene | Forward | Reverse |
| ANP | GAGAGACGGCAGTGCTTCTA<br>GGC | CGTGACACACCACAAGGGCTTAG<br>G |
| β-MHC | GAC GAG GCA GAG CAG ATC<br>GC | GGG CTT CAC AGG CAT CCT TAG<br>GG |
| NOX2 | TGC CTC CAT TCT CAA GTC | ATT CAT CCC AGC CAG TAA |
| p22phox | CAGGTGTGCTCATCTGTC | CTTCACCACAGAGGTCAGGT |
| β-actin | AGAGGGAAATGGTGCCTG AC | CAATAGTGATGACCTGGCCGT |
| Col1a1 | AATGGTGCTCCTGGTATTGC | GGCACCAGTGTCTCCTTTGT |
| TGF-β 1 | GACGTCACTGGAGTTGTACG<br>G | GCTGAATCGAAAGCCCTGT |
| F4/80 | CTTTGGCTATGGGCTTCCAGT<br>C | GCAAGGAGGACAGAGTTTATCGTG |
| TNF-α | CAGGCGGTGCCTATGTCTCA | GGCTACAGGCTTGTCACTCG |
| 18S | CGGCTACCACATCCAAGGAA | GCTGGAATTACCGCGGCT |
| Mouse primers for ddPCR |  |  |
| Gene | Forward/Reverse | Probe |
| mtDNA (ND1) | CCATTTGCAGACGCCATAAA<br>/GAGTGATAGGGTAGGTGCAA<br>TAA | ACGCCCTTTAACAACCTCT |
| ncDNA (Ago1) | GCGGTATTTCCTCTTCATCTG<br>T/GTGAGGGAACATGAGCAAT<br>CT | GTGTGTATCTCAGGCC |
| Antibodies |  |  |
| Reagent | Source | Identifier |
| 4HNE | Abcam | ab46545 |
| NQO1 | Abcam | ab13243 |
| HO-1 | Abcam | ab34173 |
| PGC1α | CST | 4259 |
| OXYPHOS<br>Rodent<br>WB antibody<br>cocktail | Abcam | ab110413 |
| TOM20 | CST | 42406 |
| Goat anti-rabbit IgG, HRP-linked Antibody | CST | 7074S |
| Goat anti-rabbit IgG H&L (HRP) | Abcam | ab6721 |
| Donkey anti-mouse IgG | Millipore | AP129P |
| **Antibodies for flow cytometry** |  |  |
| Reagent | Source | Identifier |
| LIVE/DEAD (FITC) | Invitrogen | L34969 |
| CD45 (BV421) | Biolegend | 147719 |
| CD3 (AF700) | Biolegend | 100215 |
| CD4 (BV510) | Biolegend | 100559 |
| IL-17A (PE) | Biolegend | 506903 |
| IFN-γ (AF647) | Biolegend | 505816 |
| **ELISA kits** |  |  |
| Mouse IL-12p40 | Abcam | ab236717 |
| Mouse IL-12p19 | R&D | M2300 |
| Mouse IL-12p70 | Protentech | KE10014 |
| Mouse IL-17A | R&D | M1700 |
| Human IL-12p40 | R&D | DP400 |
| **Other assay kits** |  |  |
| ATP Assay Kit-Luminescence | Dojindo | A550 |
| Mouse Cytokine array XL | R&D | ARY028 |
| **Antibodies for Neutralization** |  |  |
| Anti-mouse IL-17A | BioXcell | BE0173 |
| Anti-mouse IgG1 isotype control | BioXcell | BE0083 |

### 1 Abbreviations:

ANP, atrial natriuretic peptide; β-MHC, β-myosin heavy chain; NOX2, NADPH oxidase 2; p22phox, phagocyte oxidase subunit p22phox; β-actin, beta-actin; Col1a1, collagen type I alpha 1 chain; TGF-β1, transforming growth factor-beta 1; F4/80, macrophage marker F4/80; TNF-α, tumor necrosis factor-alpha; 18S, 18S ribosomal RNA; mtDNA, mitochondrial DNA; ND1, NADH dehydrogenase 1; ncDNA, nuclear DNA; Ago1, Argonaute 1; 4-HNE, 4-hydroxynonenal; NQO1, NAD(P)H quinone dehydrogenase 1; HO-1, heme oxygenase-1; PGC-1α, peroxisome proliferator-activated receptor gamma coactivator 1-alpha; OXPHOS, oxidative phosphorylation; TOM20, translocase of outer mitochondrial membrane 20; IgG, immunoglobulin G; HRP, horseradish peroxidase; H&L, heavy and light chains; CD45, cluster of differentiation 45; CD3, cluster of differentiation 3; CD4, cluster of differentiation 4; IL-17A, interleukin-17A; IFN-γ, interferon-gamma; ATP, adenosine triphosphate.

**Table S1. Echocardiography data of sham and 5/6Nx mice 16 weeks after last surgery**

|  | Sham (n=11) | 5/6Nx(n=18) | *P* |
| --- | --- | --- | --- |
| **HR, /min** | 704.4 ± 19.0 | 698.2 ± 17.4 | 0.381 |
| **LVEF, %** | 83.4 ± 3.91 | 81.2 ± 4.30 | 0.168 |
| **FS, %** | 68.2 ± 4.45 | 65.5 ± 4.83 | 0.985 |
| **LVDd, mm** | 3.06 ± 0.163 | 2.78 ± 0.220 | 0.002 |
| **LVDs, mm** | 0.973 ± 0.149 | 0.961 ± 0.165 | 0.850 |
| **IVST, mm** | 0.791 ± 0.054 | 1.04 ± 0.120 | <0.001 |
| **PLVW, mm** | 0.727 ± 0.047 | 1.05 ± 0.130 | <0.001 |

**Abbreviations:** HR, heart rate; LVEF, left ventricular ejection fraction; FS, fractional shortening; LVDd, left ventricular end-diastolic diameter; LVDs, left ventricular end-systolic diameter; IVST, interventricular septal thickness; PLVW, posterior left ventricular wall thickness

**Table S2. Suppressed mitochondrial- and energy metabolism–related KEGG pathways in cardiomyocytes under chronic kidney disease conditions**

| Direction | Pathway | NES | padj | GeneSetSize | LeadingEdge_n |
| --- | --- | --- | --- | --- | --- |
| Down | OXIDATIVE PHOSPHORYLATION | -2.82868415905328 | 2.12393982935147E-17 | 120 | 79 |
| Down | PARKINSONS DISEASE | -2.72735235113784 | 2.21371835358872E-15 | 114 | 75 |
| Down | CITRATE CYCLE TCA CYCLE | -2.20303491309799 | 0.000257898925653183 | 27 | 15 |
| Down | ALZHEIMER’S DISEASE | -2.04122138877409 | 5.63529695132951E-06 | 153 | 73 |
| Down | HUNTINGTON’S DISEASE | -2.03949746621528 | 5.63529695132951E-06 | 161 | 73 |
| Down | CARDIAC MUSCLE CONTRACTION | -1.98138391024786 | 0.000894553281982209 | 70 | 38 |
| Down | PANTOTHENATE AND COA BIOSYNTHESIS | -1.85596464644585 | 0.0494002057944382 | 14 | 8 |
| Down | RIBOSOME | -1.79402696724999 | 0.00817310057214486 | 79 | 34 |
| Down | SPLICEOSOME | -1.63114169639132 | 0.028566099131515 | 117 | 47 |
| Down | PYRUVATE METABOLISM | -1.57039300201483 | 0.176099795759358 | 35 | 11 |

**Abbreviations:** GSEA, gene set enrichment analysis; KEGG, Kyoto Encyclopedia of Genes and Genomes; NES, normalized enrichment score; padj, adjusted *P* value (false discovery rate–corrected); GeneSetSize, number of genes included in each KEGG pathway; LeadingEdge\_n, number of genes contributing to the leading-edge subset.

**Table S3. Upregulated KEGG pathways identified by gene set enrichment analysis (GSEA) in cardiomyocytes from CKD mice**

| Direction | Pathway | NES | padj | GeneSetSize | LeadingEdge_n |
| --- | --- | --- | --- | --- | --- |
| Up | CYTOKINE CYTOKINE RECEPTOR INTERACTION | 1.70964264754663 | 0.00233487054294842 | 187 | 71 |
| Up | PROTEIN EXPORT | 1.69431734532208 | 0.1479183547457 | 22 | 13 |
| Up | PYRIMIDINE METABOLISM | 1.61657544286735 | 0.0501511915603219 | 92 | 28 |
| Up | O GLYCAN BIOSYNTHESIS | 1.53538101669055 | 0.409691629955947 | 21 | 10 |
| Up | PROXIMAL TUBULE BICARBONATE RECLAMATION | 1.50291265372653 | 0.409691629955947 | 19 | 5 |
| Up | AXON GUIDANCE | 1.48886032873261 | 0.102312011845094 | 121 | 29 |
| Up | NICOTINATE AND NICOTINAMIDE METABOLISM | 1.4724725442281 | 0.409691629955947 | 23 | 7 |
| Up | HEMATOPOIETIC CELL LINEAGE | 1.46262910992283 | 0.306456305116969 | 67 | 34 |
| Up | GLYCINE SERINE AND THREONINE METABOLISM | 1.45751233690743 | 0.433768656716418 | 30 | 8 |
| Up | CYTOSOLIC DNA SENSING PATHWAY | 1.44757986217669 | 0.346490932321342 | 39 | 12 |

**Abbreviations:** GSEA, gene set enrichment analysis; KEGG, Kyoto Encyclopedia of Genes and Genomes; NES, normalized enrichment score;

padj, adjusted P value (false discovery rate–corrected); GeneSetSize, number of genes included in each KEGG pathway; LeadingEdge\_n,

number of genes contributing to the leading-edge subset.

**Table S4. Echocardiography data of sham, 5/6Nx and 5/6Nx mice 16 weeks after last surgery**

|  | Sham (n=12) | 5/6Nx (n=11) | 5/6Nx+anti-IL-17A (n=10) |
| --- | --- | --- | --- |
| **HR, /min** | 694.5 ± 14.8 | 697.5 ± 10.6 | 692.4 ± 11.1 |
| **LVEF, %** | 80.5 ± 3.37 | 77.4 ± 4.41 | 79.1 ± 2.72 |
| **FS, %** | 64.6 ± 3.66 | 61.0 ± 4.67 | 62.8 ± 3.17 |
| **LVDD, mm** | 2.90 ± 0.95 | 2.71 ± 0.15* | 2.73 ± 0.26 |
| **LVDs, mm** | 1.03 ± 0.106 | 1.05 ± 0.11 | 1.01 ± 0.074 |
| **IVST, mm** | 0.72 ± 0.072 | 1.08 ± 0.12* | 1.00 ± 0.12 |
| **PLVW, mm** | 0.74 ± 0.100 | 1.07 ± 0.12* | 1.00 ± 0.082 |

**Abbreviations:** HR, heart rate; LVEF, left ventricular ejection fraction; FS, fractional shortening;

LVDD, left ventricular end-diastolic diameter; LVDs, left ventricular end-systolic diameter; IVST,

interventricular septal thickness; PLVW, posterior left ventricular wall thickness. \* $P < 0.05$ vs. sham.

**Table S5. Patient clinical characteristics**

|  | All patients (n = 126) | Non-CKD (n = 27) | CKD (n = 99) | *P* |
| --- | --- | --- | --- | --- |
| Age, years | 71.6 ± 10.4 | 62.9 ± 12.09 | 74.1 ± 8.22 | <0.001 |
| Male, n (%) | 68 (54.0) | 15 (57) | 53 (54) | 0.85 |
| BMI, kg/m<sup>2</sup> | 23.7 ± 3.88 | 24.5 ± 3.86 | 23.4 ± 3.87 | 0.19 |
| Ischemic cardiomyopathy, n (%) | 18 (21) | 8 (19) | 10 (24) | 0.56 |
| Atrial fibrillation, n (%) | 50 (40) | 13 (48) | 37 (37) | 0.49 |
| Hypertension, n (%) | 83 (66) | 16 (59) | 67 (68) | 0.29 |
| Diabetes mellitus, n (%) | 47 (37) | 9 (33) | 38 (38) | 0.66 |
| Dyslipidemia, n (%) | 81 (64) | 19 (70) | 62 (63) | 0.51 |
| Hemoglobin, g/dL | 12.8 ± 2.14 | 14.1 ± 1.89 | 12.4 ± 2.08 | <0.001 |
| Serum albumin, g/dL | 3.97 ± 0.43 | 4.13 ± 0.38 | 3.92 ± 0.45 | 0.026 |
| HbA1c, % | 6.24 ± 0.84 | 6.25 ± 0.98 | 6.25 ± 0.81 | 0.99 |
| BNP, pg/mL | 56.0 (24.8–129.9) | 36.7 (19.2–62.5) | 60.2 (32.5–154.1) | 0.080 |
| hs-TnT, ng/mL | 0.025 (0.0097–0.023) | 0.0099 (0.0069–0.015) | 0.016 (0.010–0.026) | < 0.001 |
| IL-12p40, pg/mL | 116.6 (85.6–158.3) | 96.9 (72.2–123.0) | 122.7 (87.2–169.0) | 0.031 |
| Creatinine, mg/dL | 0.93 ± 0.25 | 0.83 ± 0.22 | 1.04 ± 0.24 | < 0.001 |
| eGFR, mL/min/1.73 m<sup>2</sup> | 52.3 ± 17.0 | 56.6 ± 16.4 | 47.9 ± 16.7 | 0.017 |
| LVEF, % | 58.7 ± 11.3 | 58.8 ± 10.7 | 58.7 ± 11.5 | 0.96 |
| LVDd, mm | 44.9 ± 6.83 | 46.1 ± 6.58 | 44.6 ± 6.89 | 0.28 |
| IVSTd, mm | 10.5 ± 1.92 | 9.96 ± 1.39 | 10.71 ± 2.02 | 0.031 |
| PLVWd, mm | 10.03 ± 1.36 | 10.00 ± 1.10 | 10.04 ± 1.43 | 0.88 |
| LVMi, g/m<sup>2</sup> | 100.5 ± 25.7 | 95.6 ± 19.1 | 101.9 ± 27.2 | 0.26 |
| E/e' | 13.6 ± 6.12 | 11.1 ± 3.50 | 14.3 ± 6.51 | 0.018 |
| Medications |  |  |  |  |
| Loop diuretics, n (%) | 25 (20) | 3 (11) | 22 (23) | 0.28 |
| MRA, n (%) | 13 (11) | 3 (11) | 10 (10) | 0.92 |
| Tolvaptan, n (%) | 4 (3) | 0 (0) | 4 (4) | 0.58 |
| ARB, n (%) | 48 (39) | 6 (22) | 42 (44) | 0.047 |
| β-blockers, n (%) | 57 (46) | 13 (48) | 44 (46) | 0.83 |
| CCB, n (%) | 61 (50) | 13 (48) | 48 (50) | 0.87 |
| Statins, n (%) | 21 (49) | 19 (45) | 40 (47) | 0.74 |

##### Abbreviations:

CKD, chronic kidney disease; BMI, body mass index; BNP, B-type natriuretic peptide; hs-TnT, high-sensitivity troponin T; IL-12p40, interleukin-12p40; eGFR, estimated glomerular filtration rate; LVEF, left ventricular ejection fraction; LVDd, left ventricular end-diastolic diameter; IVSTd, interventricular septal thickness at end-diastole; PLVWd, posterior left ventricular wall thickness at end-diastole; LVMi, left ventricular mass index; MRA, mineralocorticoid receptor antagonist; ARB, angiotensin II receptor blocker; CCB, calcium channel blocker.

#### Supplemental Figure Legends

##### **Figure S1. Histological and molecular characterization of renal injury in 5/6 nephrectomized mice.**

(A) Representative periodic acid–Schiff (PAS) staining (upper panels) and Picrosirius Red staining (lower panels) of kidney sections from sham-operated and 5/6 nephrectomized (5/6Nx) mice. Scale bars = 50 μm.

(B) Renal mRNA expression of *Col1a1*, αSMA, F4/80, and TNFα, determined by quantitative real-time PCR. Data are presented as mean ± SD. Statistical significance was assessed using an unpaired Student's *t* test.

##### **Figure S2. Reciprocal relationship between oxidative phosphorylation and cytokine signaling in cardiomyocytes from 5/6 nephrectomized mice.**

(A) Heatmap showing the expression of leading-edge genes in the KEGG *cytokine–cytokine receptor interaction* pathway identified by gene set enrichment analysis of RamDA-seq data obtained from isolated cardiomyocytes of sham and 5/6 nephrectomized (5/6Nx) mice. Gene expression is displayed as Z-scores.

(B) Correlation between the leading-edge gene-based oxidative phosphorylation (OXPHOS) pathway score and cytokine–cytokine receptor interaction pathway score in individual cardiomyocyte transcriptomes. Pathway activity was quantified using leading-edge gene-based pathway scores. Correlation was assessed using Spearman's rank correlation coefficient.

**Figure S3. Progression of renal dysfunction and left ventricular hypertrophy after 5/6 nephrectomy.**

(A) Serum creatinine and blood urea nitrogen (BUN) levels at 12 and 16 weeks after 5/6 nephrectomy (5/6Nx).

(B) Left ventricular ejection fraction (LVEF) and left ventricular weight normalized to tibial length (LV/TL) at 12 and 16 weeks after 5/6Nx, demonstrating preserved systolic function despite progressive cardiac hypertrophy. Data are presented as mean ± SD. Statistical significance was assessed using an unpaired Student's *t* test.

**Figure S4. Re-analysis of published cardiac single-nucleus RNA sequencing datasets reveals minimal Il12b expression and no increase in Th17-like cells in 5/6 nephrectomized hearts.**

(A) UMAP visualization of cardiac immune cell populations with cluster annotation (upper panel) and feature plot showing Il12b expression (lower panel) in the re-analyzed cardiac snRNA-seq dataset.

Il12b expression was detected in only a single nucleus.

(B) Feature plots showing the Th17 signature score and the expression of *Rorc*, *Ccr6*, and *Il23r* in cardiac T cells.

(C) Proportion of Th17-like cells among cardiac T cells in ischemia–reperfusion injury (IRI), sham, and 5/6 nephrectomy (5/6Nx) datasets.

**Figure S5. Gating strategy used for flow cytometric analysis of splenic Th17 cells.**

Representative flow cytometry plots showing sequential gating of splenocytes to identify CD45<sup>+</sup>CD4<sup>+</sup> T cells and intracellular IL-17A- and IFN-γ-producing cells. Isotype controls were used to establish gating for cytokine-positive populations.

**Figure S6. Flow diagram illustrating the patient selection process for serum IL-12p40 analysis.**

Flow chart illustrating the patient selection process for serum IL-12p40 measurement according to the predefined inclusion and exclusion criteria. A control cohort with preserved renal function was selected using the same exclusion criteria.

Unedited membranes and gels of Figure 2C.

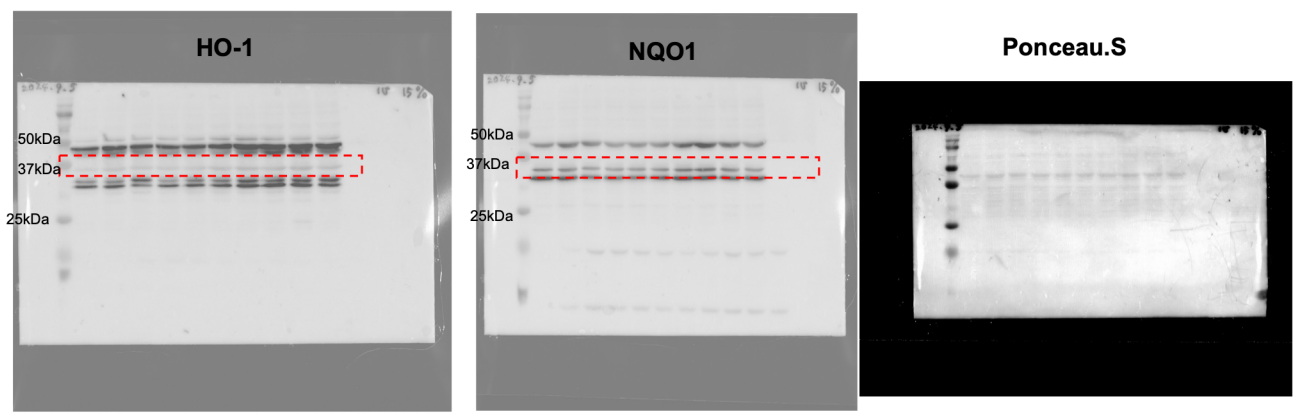

Unedited membranes and gels of Figure 3F.

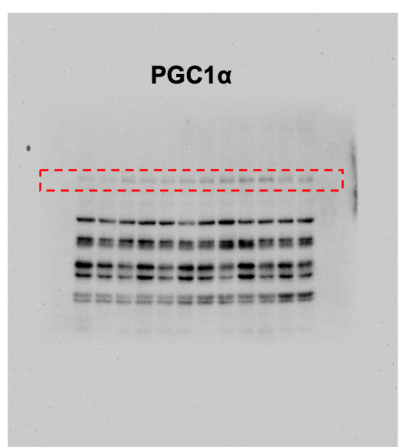

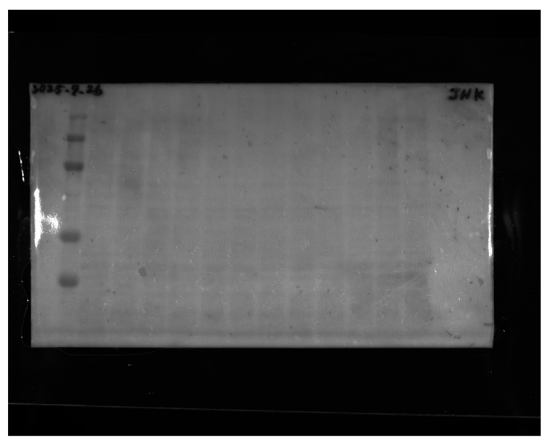

Unedited membranes and gels of Figure 3H.

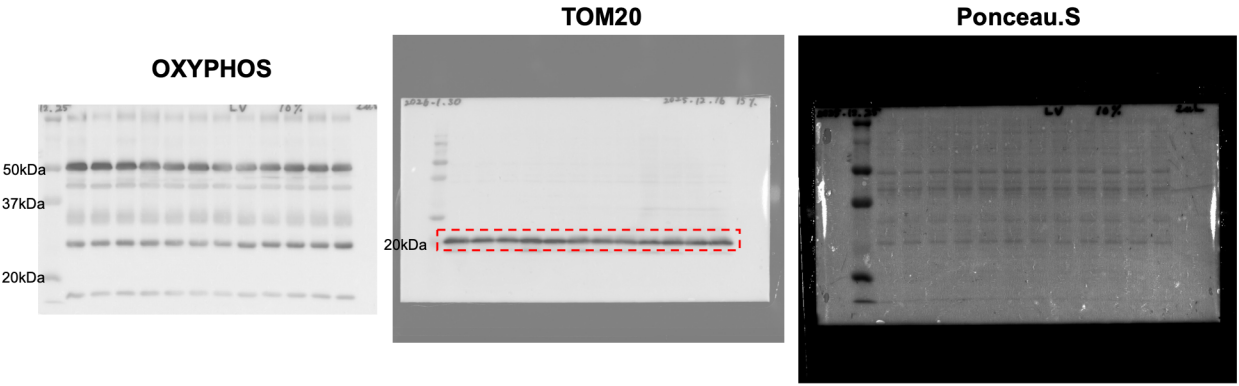

Unedited membranes and gels of Figure 5G.

OXYPHOS

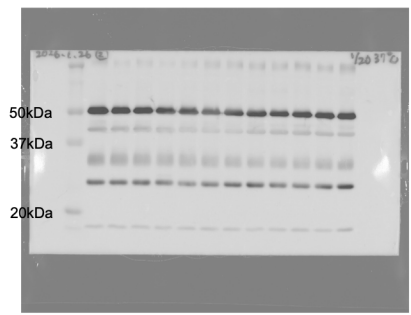

TOM20

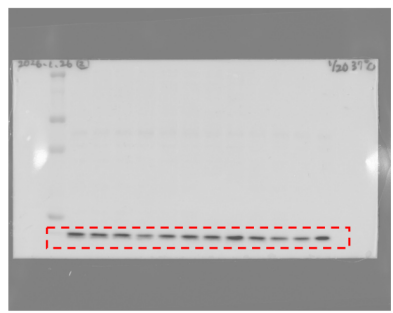

Ponceau.S

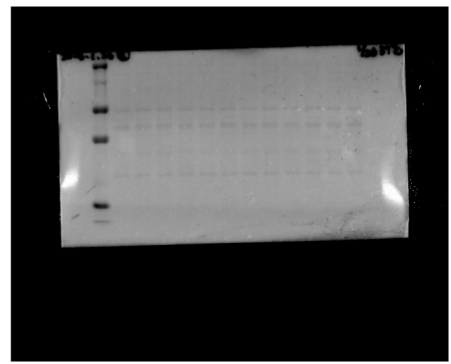
