## Supplementary figures and images for "IL-17A Mediates Cardiac Hypertrophy in Chronic Kidney Disease through the IL-23/Th17 Axis"

### Figures S1- S6

Figure S1.

A.

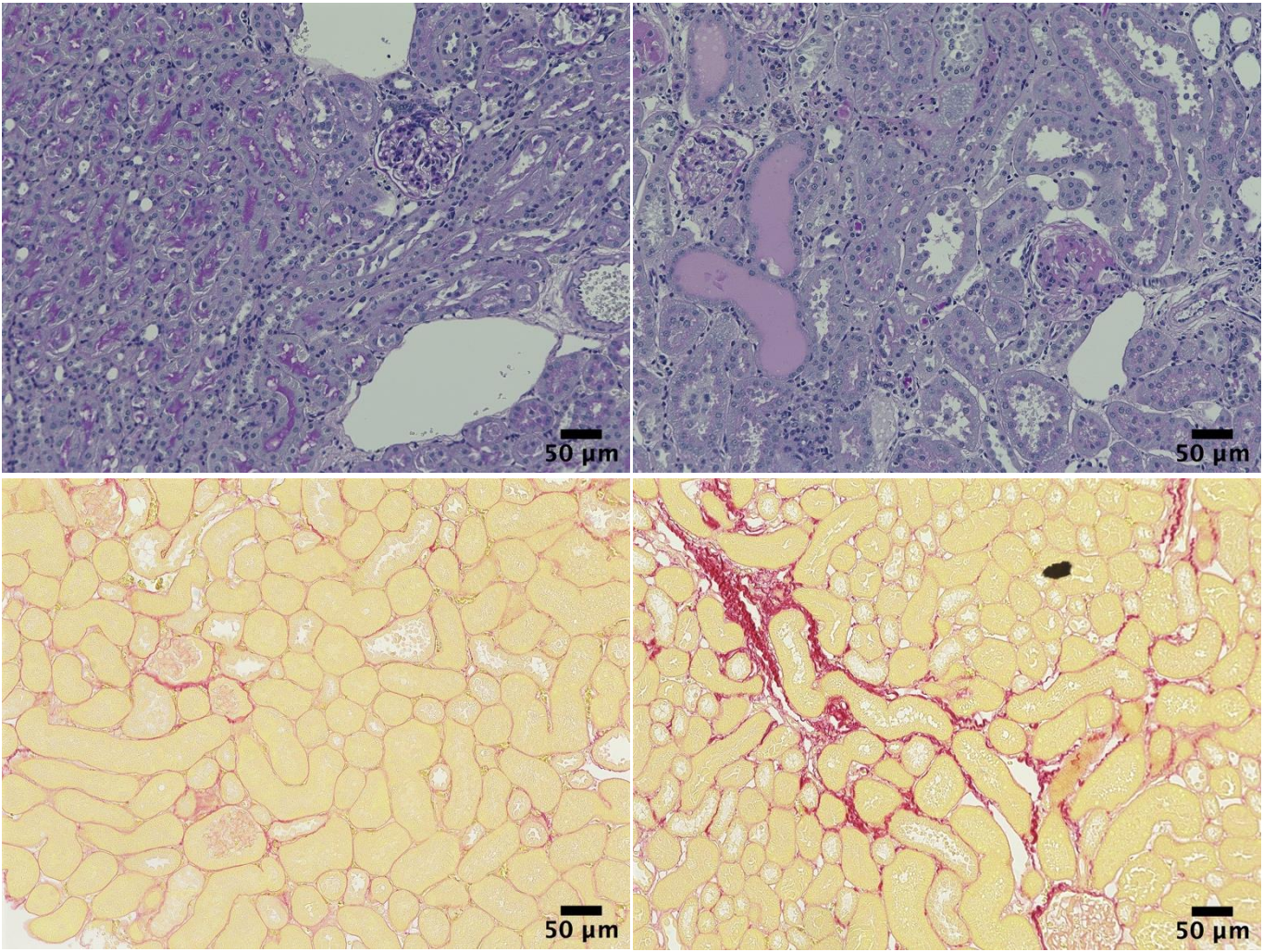

B.

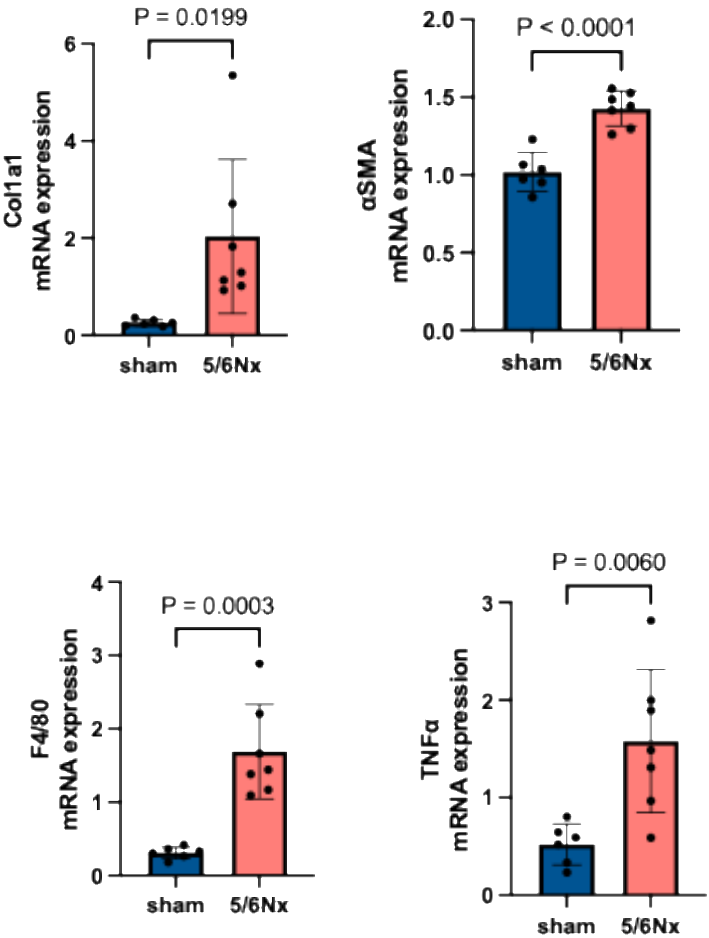

Figure S2.

A.

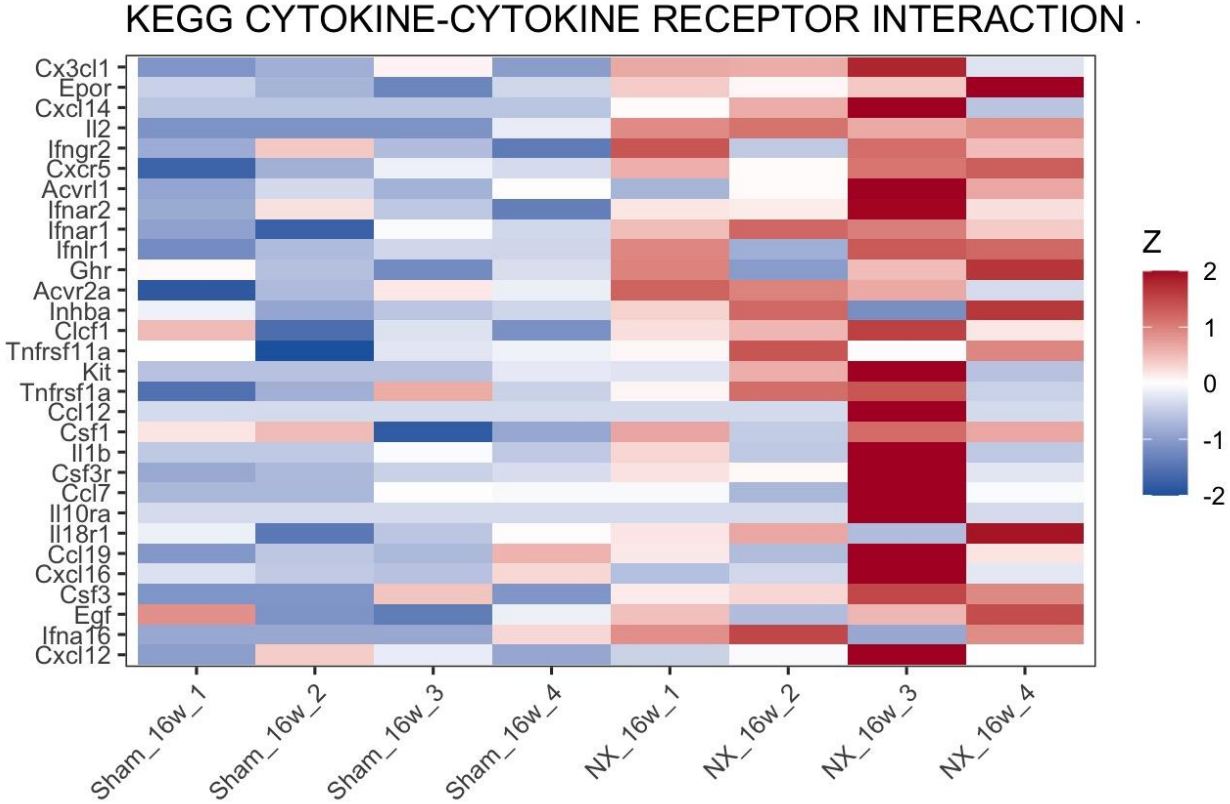

B.

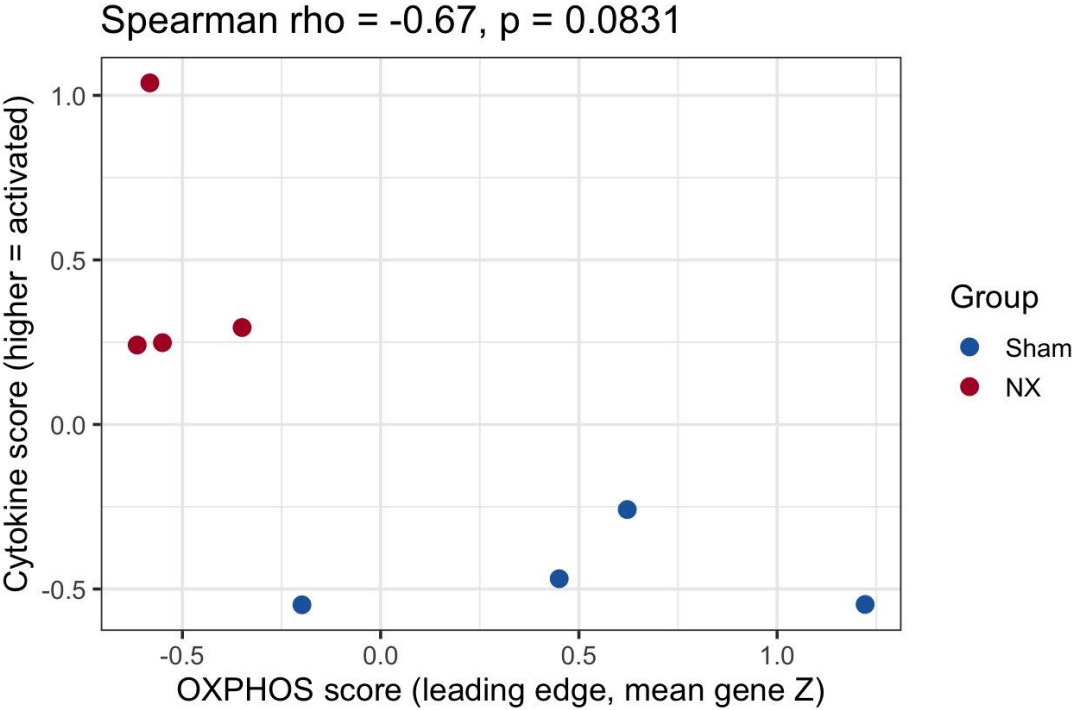

Figure S3.

A.

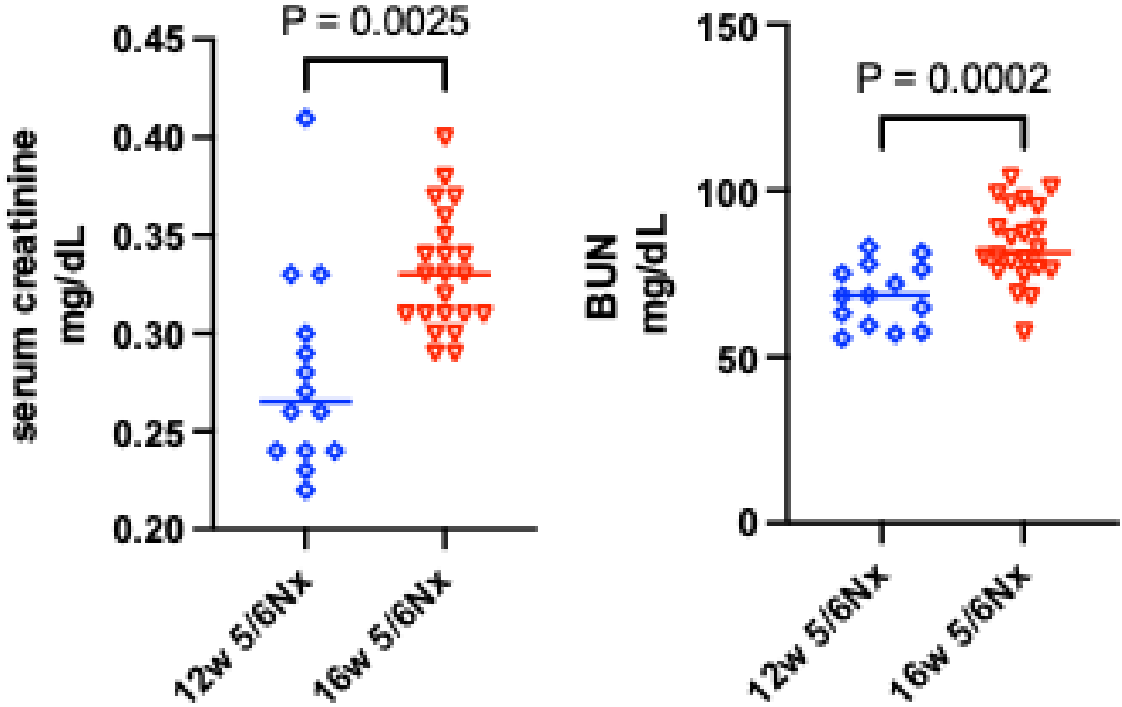

B.

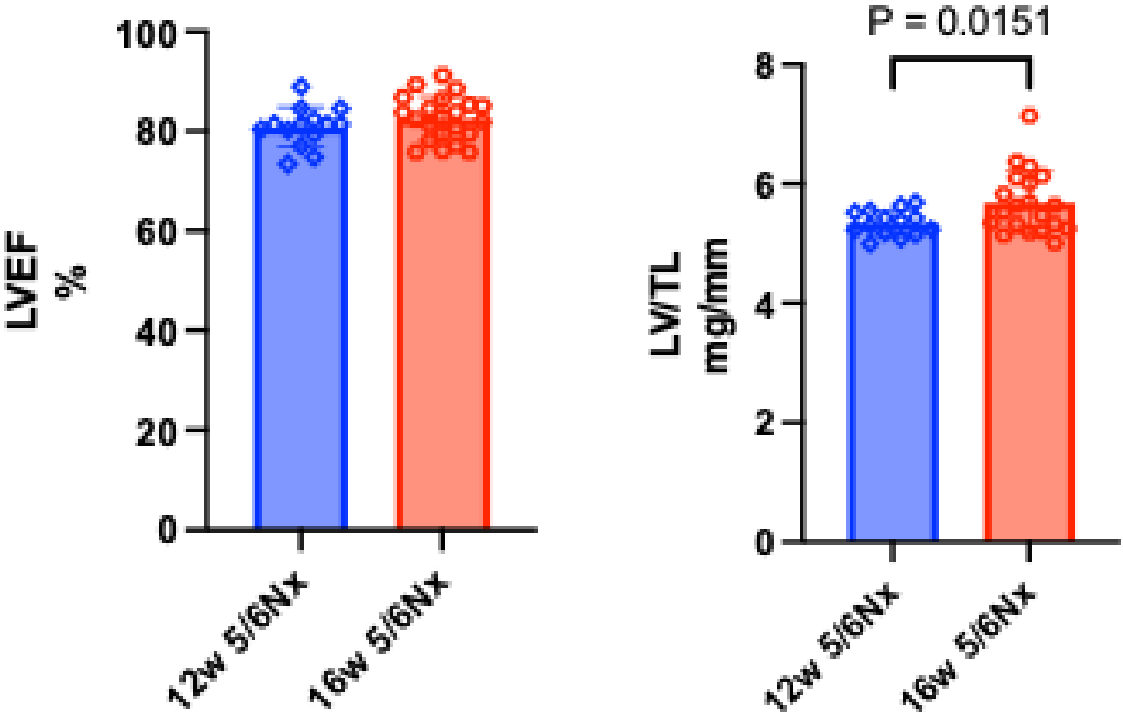

Figure S4. Reanalysis of GSE180852

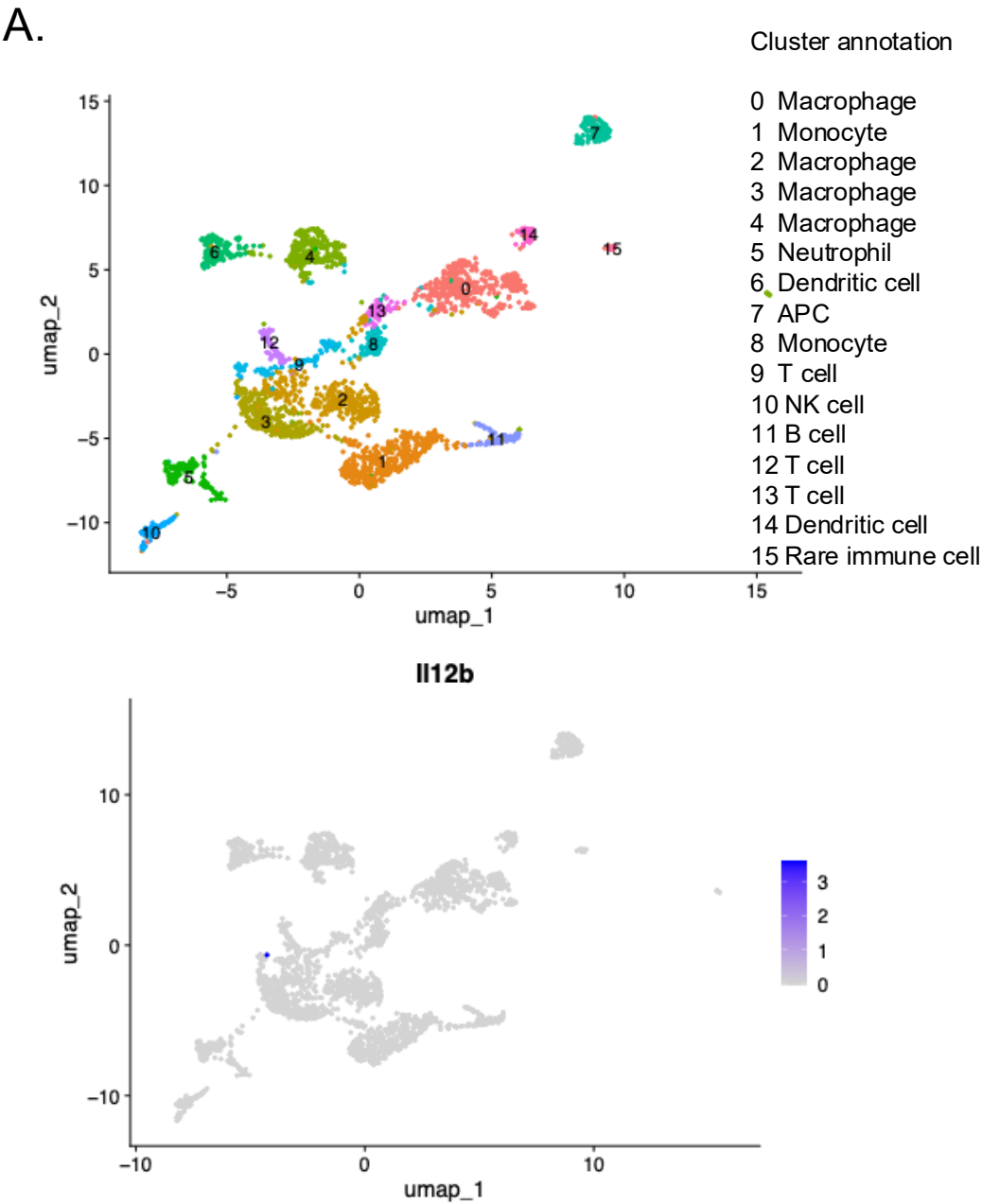

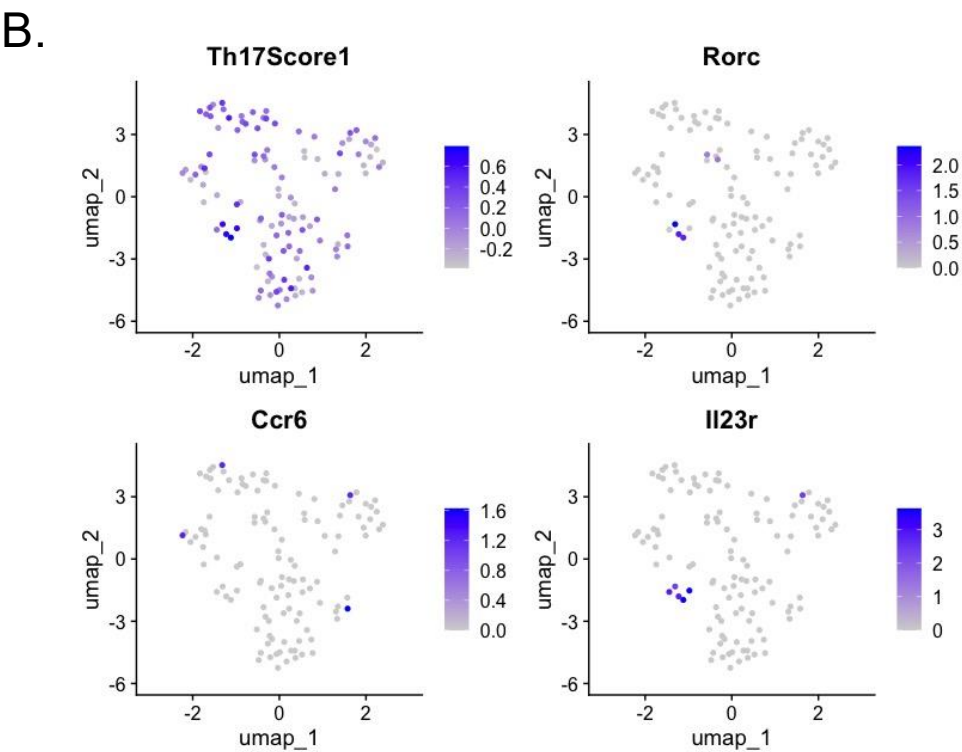

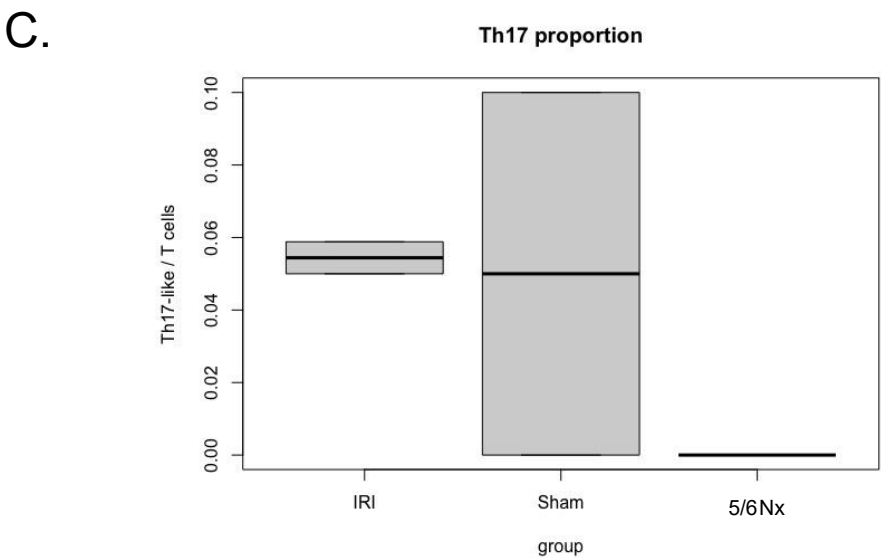

Figure S5.

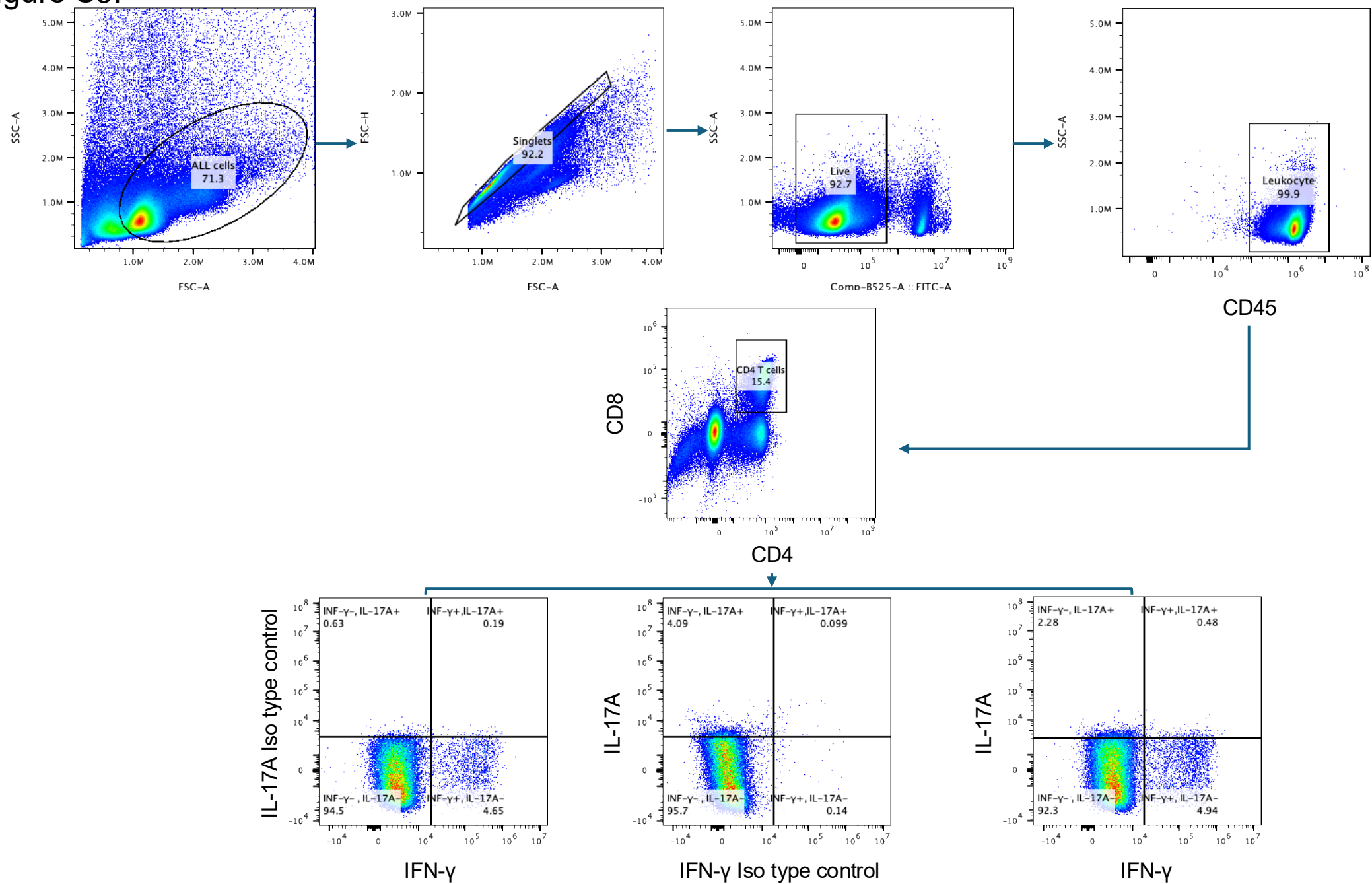

Figure S7.

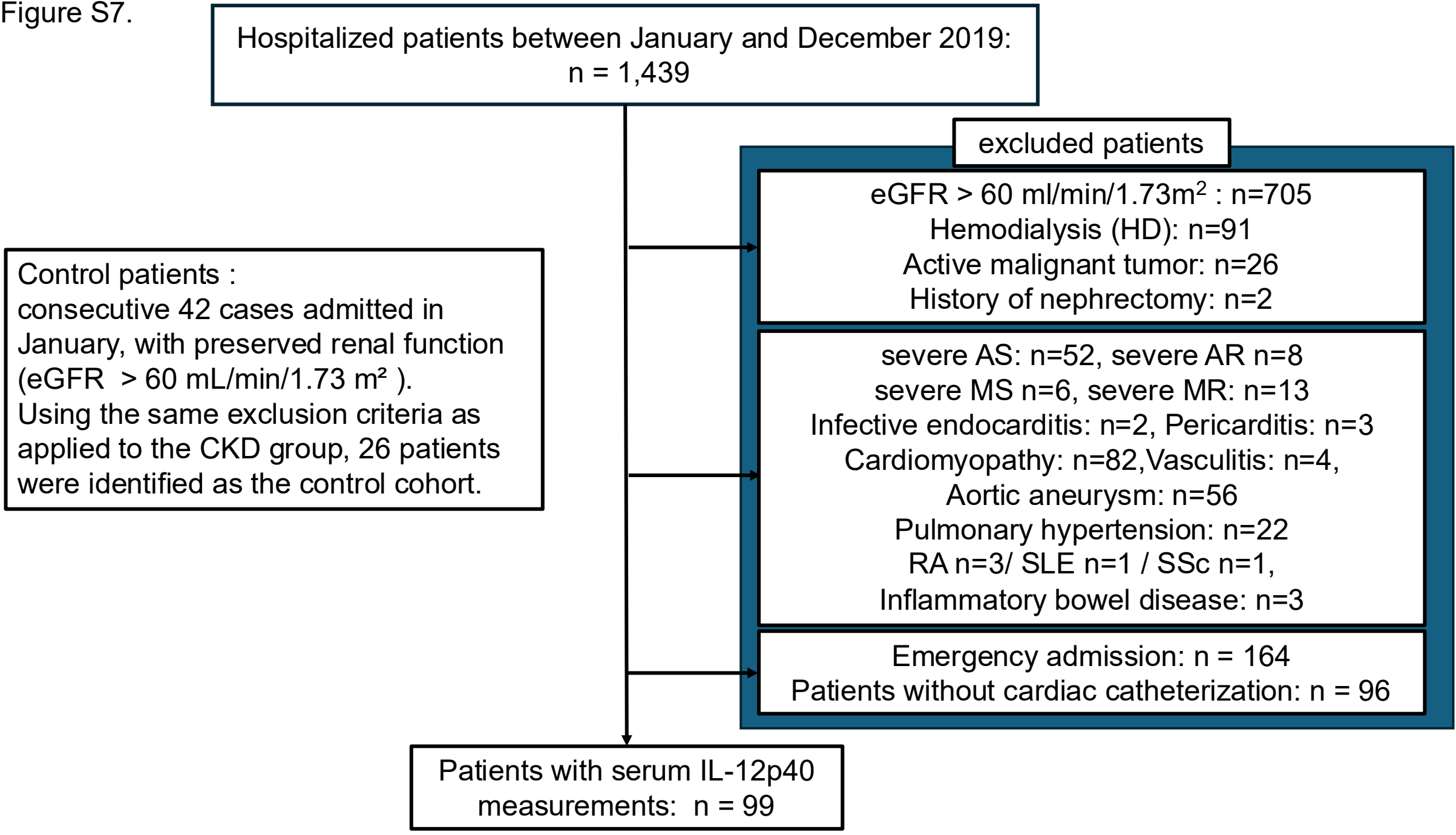
